# Temperature elevation synergises with and enhances the type-I IFN-mediated restriction of MPXV

**DOI:** 10.1101/2023.09.29.560106

**Authors:** Chris Davis, Ilaria Epifano, Kieran Dee, Steven McFarlane, Joanna K. Wojtus, Benjamin Brennan, Quan Gu, Vattipally B. Sreenu, Kyriaki Nomikou, Lily Tong, Lauren Orr, Ana Da Silva Filipe, David A. Barr, Antonia Ho, Emma C. Thomson, Chris Boutell

## Abstract

Fever is an evolutionary conserved host pro-inflammatory immune response that governs the regulation of multiple biological processes to control the outcome of infection. In January 2022, the World Health Organization (WHO) reported a global outbreak in mpox cases with a high incidence of human-to-human transmission. A frequent prodromal symptom of monkeypox virus (MPXV) infection is fever, with a febrile temperature range of 38.3 to 40.5 °C. However, the outcome of temperature elevation on MPXV infection remains poorly defined. Here, we isolated a circulating strain of MPXV from a patient who presented with fever (38.5 °C) and rash from the 2022 outbreak. Genomic sequencing identified this isolate to belong to the epidemic Clade IIb.B1. Transcriptomic analysis of infected cells demonstrated this virus to induce a strong IL6 pro-inflammatory immune response, consistent with a role for this pyrogen in the regulation of fever. We identify host-cell temperature at both physiological skin (33 °C) and clinical febrile temperatures (38.5 and 40 °C) to be a key determinant in the outcome of infection through the differential regulation of MPXV transcription and associated amplitude of host cytokine response to infection. Incubation of infected cells at 38.5 or 40 °C led to a restriction or ablation in MPXV replication, respectively. Importantly, this thermal inhibition was reversible upon temperature downshift to 37 °C without detriment to viral replication fitness. Co-stimulation of the type-I interferon (IFN) response led to a dose- and temperature-dependent inhibition in MPXV replication that restricted the re-establishment of infection upon temperature downshift and withdrawal of IFN as an immune stimulus. Our data identify febrile temperatures associated with mpox disease to be a critical component of the host pro-inflammatory immune response to infection which can synergise with the type-I IFN response to enhance the host-cell mediated restriction of MPXV.

## Introduction

MPXV belongs to the Orthopoxvirus (OPXV) genus^1^, which includes variola virus (VARV, the causative agent of smallpox) and vaccinia virus (VACV, the foundation virus of the smallpox vaccine). Since January 2022, the World Health Organization (WHO) reported a global rise in the number of confirmed mpox cases across all six WHO regions (> 90,000 confirmed cases and ≥ 150 deaths; September 2023), including previous non-endemic regions of Europe and the Americas (https://worldhealthorg.shinyapps.io/mpx_global/). While the natural zoonotic reservoir of MPXV is likely to be in small rodents^2-4^, increased human-to-human transmission has been widely reported during the 2022/23 outbreak and linked to the emergence of novel Clade IIb variants since 2016^5-7^. This has raised concern that MPXV may establish a global foothold in the human population due to waning levels of cross-protecting immunity conferred previously through smallpox vaccination^8,9^.

Human-to-human transmission of MPXV occurs through close contact with infectious skin lesions, bodily fluids, or large respiratory droplets^3,4^. The spectrum of mpox disease is often variable and ranges from self-limiting disease in healthy immunocompetent adults to fatality in 1 to 10 % of affected individuals^10^. Common prodrome symptoms include fever, lymphadenopathy, pharyngitis, headache, and myalgia, followed by rash and skin lesions on the body, face, and genitals^8,10-13^. More than 70 % of confirmed cases report fever, with a febrile temperature range of 38.3 to 40.5 °C^11,14-17^. Fever is an evolutionary conserved host response to infection and inflammation that can influence multiple cellular processes, including host immunological responses to infection^18,19^. Body temperature varies throughout the day, with age, sex, and ethnic origin being contributing factors^20,21^. Unlike heat stroke or hyperthermia, fever represents a controlled shift in body temperature activated in response to exogenous (microbial) and endogenous (host) pyrogenic factors^20-22^. Clinical febrile temperatures can vary (ΔT of 1 to 4 °C above baseline); with low (38 to 39 °C), moderate (39.1 to 40 °C), high (40.1 to 41 °C), and hyperpyrexia (> 41.1 °C) temperature ranges^21^. Experimental evidence has shown tissue temperature to play a key role during viral infection and the induction of interferon (IFN)-mediated antiviral innate immune defences to infection^23-26^. Notably, antipyretic treatment of intensive care patients infected with influenza A virus (IAV) has been linked to increased patient mortality^27-29^, suggesting a beneficial role of the fever response to protect against infection in a clinical setting.

While small animal and non-human primate (NHP) models have shown MPXV infection to induce fever^30-32^, few studies have directly examined the influence of temperature on the outcome of infection. Historical evidence (circa 1960s) has shown divergent OPXVs to have variable ceiling temperatures of pox formation in chick embryos, with mpox restriction observed at temperatures ≥ 39.5 °C^33,34^. These findings suggest that tissue temperature is likely to play a key determinant in the replication and/or immunopathology of MPXV. However, *in vitro* studies have focused on laboratory-adapted strains of VACV, which are known to carry strain-specific mutations that can influence their thermal- and/or immuno-regulatory properties^22,35-38^. Thus, the influence of temperature elevation to control the outcome of OPXV infection remains poorly defined, specifically in circulating Clades or strains of human origin. We therefore set out to investigate the net effect of temperature elevation on the replication of MPXV using a clinical isolate derived from the 2022/23 outbreak.

We isolated a circulating strain of MPXV from a febrile hospitalised patient during the 2022 outbreak, who was recruited to the International Severe Acute Respiratory and Emerging Infections Consortium Comprehensive Clinical Characterisation Collaboration (ISARIC4C) study (ISRCTN66726260). Genomic sequencing demonstrated this isolate belonged to the Clade IIb.B1 epidemic strain. We demonstrate basal tissue and host-cell temperature to be a key determinant in the outcome of MPXV infection at both physiological skin temperature (33 °C) and clinical febrile (38.5 and 40 °C) temperatures relative to core body temperature (37 °C). Incubation of infected cells at 40 °C led to an ablation in MPXV replication. We show this thermal restriction to occur via a genome wide suppression in viral transcription, with multiple viral open reading frames (ORFs) displaying temperature-dependent profiles of differential gene expression (DEG). Importantly, we show that the thermal restriction of MPXV is reversible upon temperature downshift to 37 °C without impairment to viral replication fitness. Co-stimulation of the type-I IFN response led to a dose- and temperature-dependent IFN-mediated restriction in MPXV replication that attenuated the re-establishment of infection upon thermal downshift. Our data identify a cooperative and synergistic role for temperature elevation to enhance the type-I IFN-mediated restriction of MPXV. Findings pertinent to the immunological regulation of many clinical pathogens that induce a fever response.

## Results

### Temperature elevation inhibits MPXV replication in a cell-type dependent manner

We isolated a clinical strain of MPXV from a PCR positive patient presenting with fever (38.5 °C), cough, myalgia, and rash. Illumina sequencing of inactivated clinical swabs (CVR_MPXV1a) and infectious cell culture supernatant derived from primary amplification (MPXV CVR_S1) identified an identical MPXV genome (accession number ON808413) (Fig 1A). Phylogenetic analysis identified this virus to belong to the MPXV Clade IIb.B1 lineage and to contain 67 single nucleotide polymorphisms (SNPs) relative to the MPXV Clade II reference strain (NC_063383). SNPs were located across the genome and present in both core and accessory ORFs. Consistent with previous MPXV Clade IIb genomic analysis^7,39,40^, 91% of the SNPs (61 out of 67 nucleotides) were consistent with APOBEC deamination (G to A or C to T substitutions; Fig 1B).

**Fig 1.**
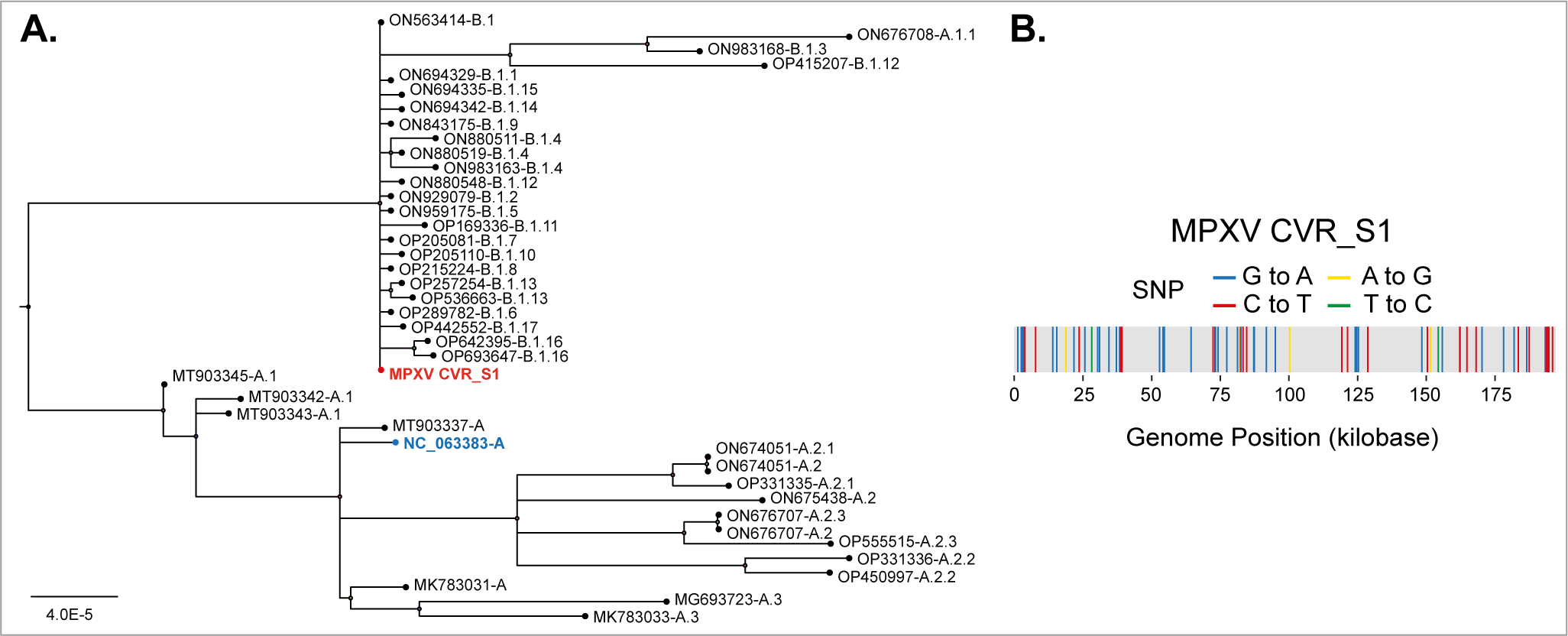
Clinical MPXV isolation and genotyping. Skin swabs obtained from a hospitalised MPXV PCR-positive patient presenting with fever and rash during the 2022 mpox outbreak were used to isolate and amplify infectious virus. Clinical swabs (CVR_MPXV1a) and supernatant derived from primary amplification (MPXV CVR_S1) were subjected to Illumina sequencing and confirmed to be identical (accession number ON808413). (A) Phylogenetic tree highlighting the evolutionary and taxonomic relationship of MPXV CVR_S1 (red text) to other sequenced MPXV Clade IIa and IIb strains (as indicated). (B) Schematic diagram highlighting the position and distribution of the 67 single nucleotide polymorphisms (SNPs; supplemental S1 data) identified within the MPXV CVR_S1 genome relative to the MPXV Clade IIa reference strain ([NC_063383-A](https://www.ncbi.nlm.nih.gov/nuccore/NC_063383); blue text) used for genome annotation throughout the study. Blue lines (guanine to adenine, G to A), red lines (cytosine to thymine, C to T), yellow lines (adenine to guanine, A to G), green lines (thymine to cytosine, T to C).

As MPXV CVR_S1 (here after referred to as MPXV) was isolated from a patient with fever, we next investigated the influence of tissue temperature on MPXV replication in skin epithelium. Human keratinocytes were differentiated into a pseudostratified epithelium under air liquid interface (ALI). Tissues were mock treated or infected with MPXV (10^2^ or 10^3^ plaque-forming units (PFU)/tissue) at 37 °C for 1 h prior to continued incubation at 33, 37, or 40 °C (representative of skin, core, and maximum clinical febrile temperature range, respectively) for 72 h. Haematoxylin and eosin (H&E) staining of mock treated tissue sections demonstrated no significant difference in skin epithelium thickness, morphology, or integrity over the temperature range of analysis (Fig 2A, B; Fig S1). As expected, immunohistochemistry (IHC) staining of tissue sections identified MPXV virion antigen expression to increase in a multiplicity of infection (MOI) dependent manner at 33 and 37 °C (Fig 2A; Fig S1). Notably, discrete foci of infection could be observed at both temperatures, indicative of MPXV intraepithelial propagation and spread at 72 h (Fig 2A; 10^2^ PFU/tissue). Quantitation of IHC stained tissue sections demonstrated the relative levels of MPXV infection to be dependent on incubation temperature, with the highest levels of replication observed at 37 °C (Fig 2C, D). Importantly, a significant reduction in MPXV antigen staining could be observed at both physiological skin (37 *vs*. 33 °C) and febrile (37 *vs.* 40 °C) incubation temperatures, with MPXV staining at 40 °C close to background levels observed in mock treated samples (Fig 2A, C, D). Together, these data identify tissue temperature to play a key determinant in the outcome of MPXV replication at both physiological and febrile temperature ranges associated with clinical infection.

**Fig 2.**
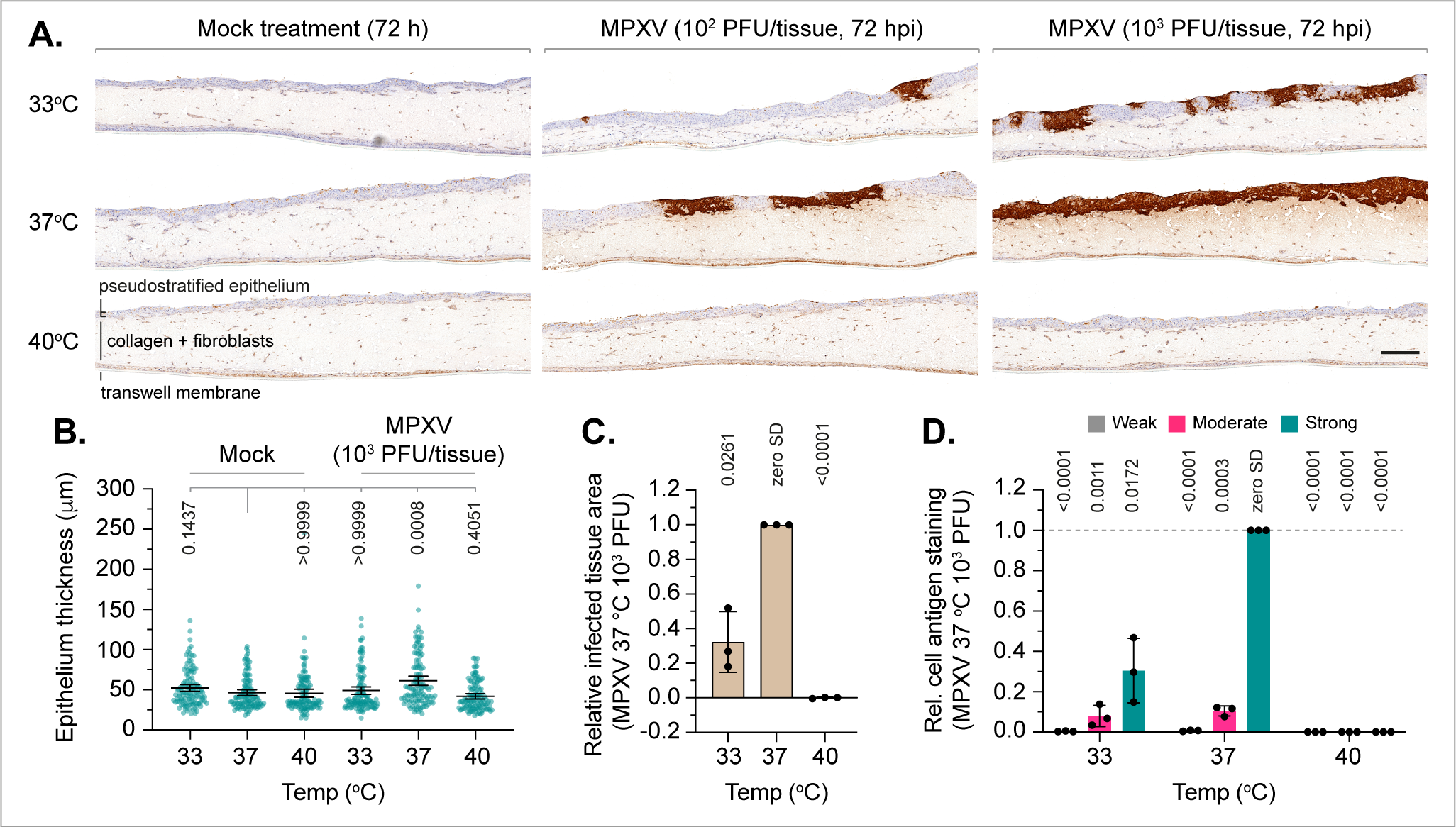
Tissue temperature influences the outcome of MPXV replication in skin epithelium. Human keratinocytes were differentiated into pseudostratified skin epithelium under air liquid interface (ALI) for 12 to 14 days. Tissues were mock treated or MPXV infected (MOI 10^2^ or 10^3^ PFU/tissue) for 1 h at 37 °C prior to incubation at 33, 37, or 40 °C for 72 h (as indicated). (A) Representative immunohistochemistry (IHC) stained tissue sections counter-stained with haematoxylin. Brown highlights epithelial regions positive for MPXV virion antigen expression. Scale bar 0.25 mm. (B) Quantitation of epithelial thickness of mock treated or MPXV infected tissues (as in A). Mean and 95% CI shown. Values derived from a minimum of 40 measurements per tissue; *p*-values shown, Kruskal-Wallis one-way ANOVA. (C) Quantitation of MPXV infected tissue area. Values normalised to infected tissues incubated at 37 °C. Mean and SD shown. (D) Proportion of weak, moderate, and strong MPXV virion antigen positive cells. Values normalised to the proportion of strong MPXV antigen-stained cells from infected tissues incubated at 37 °C per biological experiment. (C/D) *P*-values shown, one sample two-tailed t test against a theoretical mean of 1. (A to D) *N*=3 independent biological experiments. Raw values presented in supplemental S2 data. Original tissue section scans shown in Fig S1.

To examine the influence of temperature on the replication kinetics of MPXV in more detail, we next compared the thermal sensitivity of MPXV to that of VACV (strain WR; internal positive control)^38^. Human foreskin fibroblast (HFt) cells were infected at 37 °C for 1 h prior to incubation at 33, 37, 38.5 or 40 °C (representative of skin, core, and low to moderate grade febrile temperature ranges, respectively). Infected monolayers were fixed at 48 h and analysed for viral plaque formation by Coomassie staining. Consistent with previous results^38^, VACV demonstrated a significant decrease in plaque number, plaque size, and titre of cell-released virus (CRV) at incubation temperatures > 37 °C (Fig 3A to D). Notably, plaque formation could still be observed at 40 °C, demonstrating the inhibitory ceiling temperature for VACV had yet to be reached (Fig 3A, B)^38^. A significant reduction in plaque size and CRV titre could also be observed upon incubation at 33 °C (Fig 3C, D), demonstrating physiological temperatures below 37 °C to be inhibitory to optimal VACV replication. qPCR analysis of intracellular viral DNA (vDNA) levels harvested at 2 h post-infection (hpi; 1 h post-temperature shift) demonstrated the thermal restriction of VACV to occur independently of a block to virus entry (Fig 3E). Moreover, no significant difference in cell number, cell viability, or puromycin-induced cell death (positive control) was detected in mock treated cells at 40 °C relative to incubation at 37 °C (Fig S2A, B), confirming cell monolayer health and responsiveness to stimuli at temperatures > 37 °C. These data validate our model system and corroborate VACV to be sensitive to thermal restriction at temperatures ≥ 38.5 °C^38^. Analogous infections demonstrated MPXV plaque formation and plaque size to be highly sensitive to alterations in incubation temperature, with reduced numbers of plaques observed at 33 °C, reduced plaque size at 38.5 °C, and a complete inhibition in viral plaque formation at 40 °C (Fig 3F to H). Significantly lower titres of CRV were also detected at 33, 38.5, and 40 °C relative to incubation at 37 °C independently of a block to virus entry (Fig 3I, J). To establish whether the temperature-dependent restriction observed in OPXV replication was a consequence of a failure of HFt cells to support DNA virus replication, we examined the plaque formation of herpes simplex virus 1 (HSV-1, strain 17syn+) over an equivalent temperature range. Only a modest restriction in plaque count was observed at 40 °C (Fig S2C), demonstrating HFt cells to support viral DNA replication over a wide range of incubation temperatures. Thus, we identify a low-passage MPXV clinical isolate to be highly sensitive to thermal restriction at both physiological (33 °C) and febrile (38.5 or 40 °C) temperatures relative to incubation at core body temperature (37 °C).

**Fig 3.**
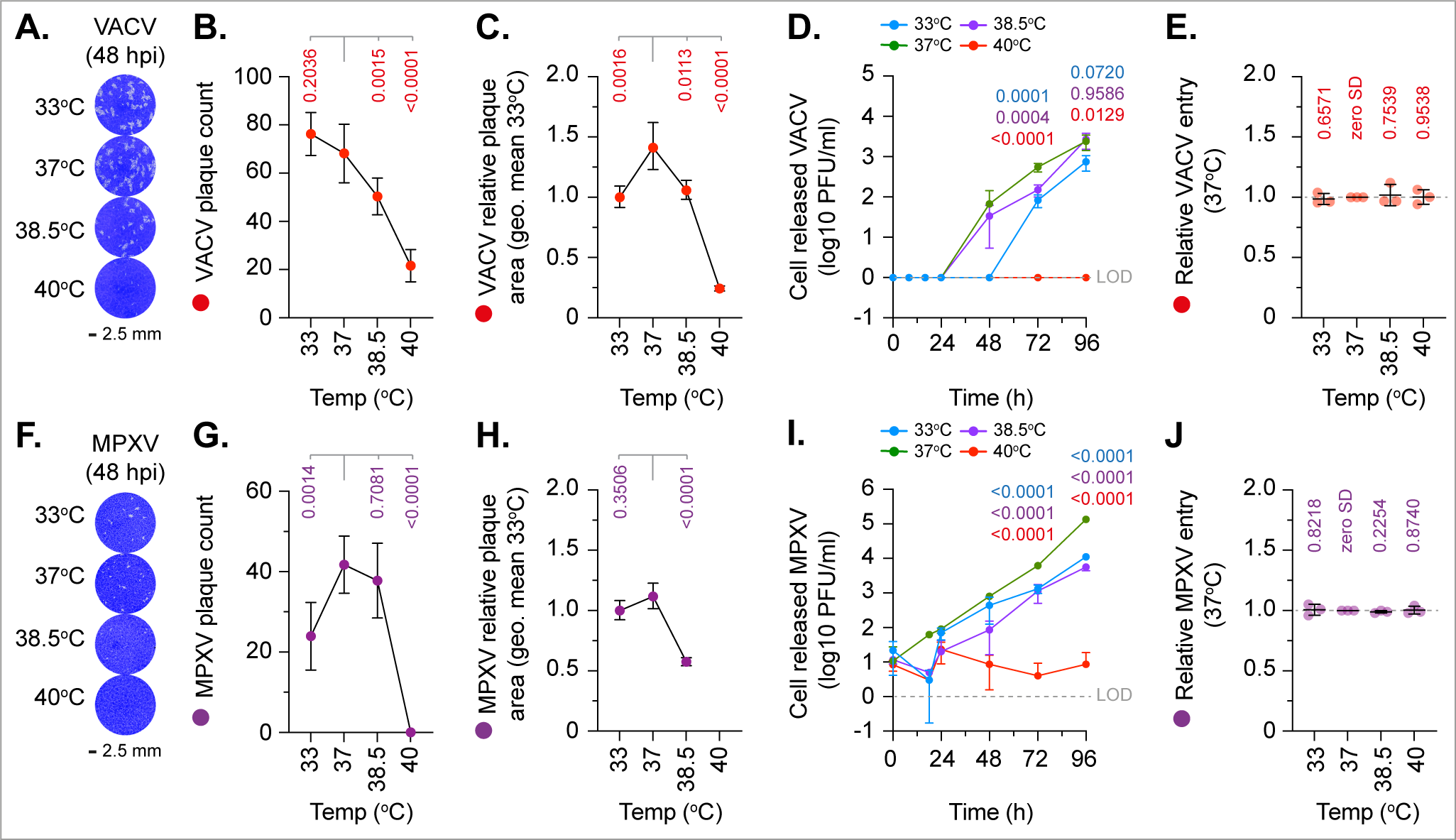
Temperature elevation inhibits MPXV replication. Human foreskin fibroblast (HFt) cells were infected with VACV (MOI 0.0005 or 0.1 PFU/cell, A to D or E, respectively; top panels) or MPXV (MOI 0.001 or 0.1 PFU/cell, F to I or J, respectively; bottom panels) for 1 h at 37 °C prior to incubation at 33, 37, 38.5 or 40 °C. (A/F) Representative images of VACV or MPXV infected cell monolayers stained with Coomassie Brilliant blue at 48 h post-infection (hpi). (B/G) Quantitation of VACV or MPXV plaque counts at 48 hpi. Means and SD shown; *p*-values shown, Dunnett’s unpaired one-way ANOVA. (C/H) Quantitation of plaque diameters at 48 hpi. Values normalised to the geometric mean at 33 °C per biological experiment. Means and 95% CI shown; *p*-values shown, Kruskai-Wallis one-way ANOVA. (D/I) Quantitation (log10 PFU/ml) of cell-released virus from infected HFt cells quantified by plaque assay in Vero E6 cells. Mean and SD shown; *p*-values shown, Dunnett’s unpaired one-way ANOVA. Limit of detection (LOD) shown (dotted grey line). (E/F) qPCR quantitation of virus host-cell entry at 1 hpi. Values normalised to 37 °C per biological experiment. All data points shown; line, mean; whisker, SD; *p*-values shown, one sample two-tailed t test against a theoretical mean of 1. (A to J) *N*≥3 independent biological experiments. Raw values presented in supplemental S3 data.

As MPXV is known to infect a wide variety of cell types *in vivo*^30^, we investigated if the thermal sensitivity of either VACV or MPXV occurred in a cell-type dependent manner. Infection of human normal oral keratinocytes (NOK) and retinal pigmented epithelial (RPE) cells with VACV demonstrated an equivalent trend in thermal restriction to that observed in HFt cells (Fig 4A, B; Fig S2D). Notably, infection of *Chlorocebus sabaeus* or *Chlorocebus aethiops* (African green monkey) kidney-derived epithelial or fibroblast cells (Vero E6 and CV-1 cells, respectively) demonstrated significantly higher levels of VACV plaque formation at 33 °C relative to incubation at 37 °C (Fig 4A, B; Fig S2D). These data demonstrate that the thermal permissivity of cells to support VACV replication to be cell-type dependent. Analogous infection with MPXV demonstrated distinct cell-type dependent profiles of thermal restriction at 33 °C (HFt, RPE, and CV-1 cells) and 38.5 °C (NOK, Vero E6, and CV-1 cells) relative to incubation at 37 °C (Fig 4C, D; Fig S2E). All cell lines demonstrated a complete, or near complete (CV-1 cells), restriction in MPXV plaque formation at 40 °C (Fig 4C, D). We conclude the basal and ceiling temperatures that support or restrict MPXV replication to be cell-type dependent and distinct from laboratory-adapted VACV.

**Fig 4.**
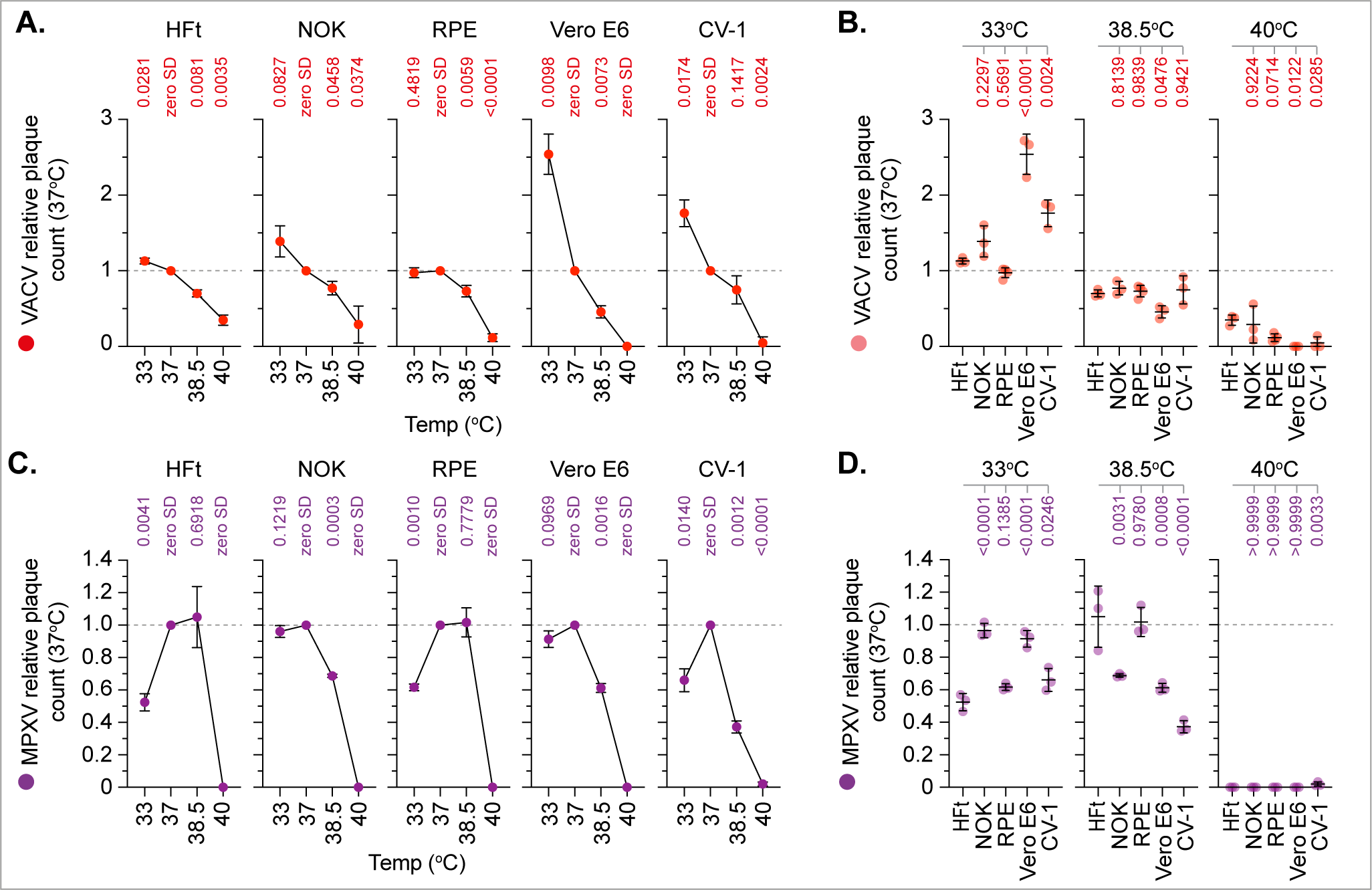
Basal host-cell temperature influences MPXV replication in a cell-type dependent manner. Confluent HFt, NOK (human normal oral keratinocyte), RPE (human retinal pigmented epithelial), Vero E6 (green monkey kidney epithelial), or CV-1 (green monkey kidney fibroblast) cell monolayers were infected with VACV (MOI 0.0005 PFU/cell; top panels) or MPXV (MOI 0.001 PFU/cell; bottom panels) for 1 h at 37 °C prior to incubation at 33, 37, 38.5 or 40 °C. Infected cell monolayers were stained with Coomassie Brilliant blue at 48 h post-infection and plaque counts quantified. Values were normalised to 37 °C per cell line per biological experiment (grey dotted line). (A/C) Normalised plaque counts per cell line over the temperature range of analysis. Means and SD shown; *p*-values shown, one sample two-tailed t test against a theoretical mean of 1. (B/D) Normalised plaque counts per temperature condition. Means and SD shown; *p*-values shown, Dunnett’s unpaired one-way ANOVA. (A to D) *N*=3 independent biological experiments. Raw values presented in supplemental S4 data.

### Basal host-cell temperature differentially regulates the outcome of MPXV transcription

To determine where the block in MPXV replication might occur, we performed RNA sequencing (RNA-Seq) on mock-treated or MPXV-infected HFt cells incubated at 33, 37, 38.5 or 40 °C (MOI 0.01 PFU/cell, 48 h). Transcriptome analysis of 28 reference genes known to be expressed across a wide range of tissues and cell-types^41-43^ demonstrated no significant difference in their relative profile of expression in mock treated cells across the temperature range of analysis (Fig S3A). Moreover, no significant difference was observed in the total number of host mapped read (MR) counts derived from mock treated samples across the temperature range (Fig S3B, grey bars). These data indicate host-cell transcription to remain largely unperturbed in mock-treated cells up to 40 °C. In contrast, a significant difference in host MR counts was observed in MPXV-infected cells incubated at 33, 37, and 38.5 °C relative to mock treatment at 37 °C (Fig S3B, green bars). No significant difference in host MR count was observed between MPXV-infected cells incubated at 40 °C and mock treatment at 37 °C. These data indicate MPXV infection to significantly influence the outcome of host transcription at incubation temperatures permissive to MPXV replication (Fig 3F to I). Investigating further, principal component analysis (PCA) of 178 MPXV ORFs identified significant variance in viral transcript expression (counts per million; CPM) levels at both physiological (33 °C) and febrile (38.5 and 40 °C) temperatures relative to incubation at core body temperature (37 °C) (Fig 5A; Fig S3C). Expression profile analysis identified infected cells incubated at 40 °C to have significantly lower levels of MPXV transcription relative to all other incubation temperatures (Fig 5B, C; Fig S3C). Notably, the overall profile of transcription remained proportionate to that observed at 37 °C, with an approximate 100-fold reduction in transcript expression per ORF at 40 °C (Fig 5B, C). 29 ORFs were identified to have counts below one CPM (Fig 5B, purple circles and text). Of these, 10 ORFs encode proteins involved in intracellular virus maturation, two ORFs that encode virus toll-like receptor (vTLR) antagonists, and three ORFs that encode subunits of the viral RNA (vRNA) polymerase complex (Fig 5D, purple text). Thus, we attribute the ablation of MPXV replication at 40 °C to suppressed levels of viral transcription that bottleneck or restrict the progress of infection. As we had also observed differences in MPXV transcription at lower incubation temperatures (Fig 5A), we next compared the transcription profiles from MPXV infected cells incubated at 33 and 38.5 °C. Relative to 37 °C (Fig 5E, solid grey line), distinct profiles of MPXV transcription could be observed between these two incubation temperatures. Notably, many ORFs displayed opposing profiles of transcription expression (Fig 5E, dotted grey ellipses). Five functionally unrelated ORFs were identified to be downregulated at 33 °C (≥ 1.5 log2 FC) relative to incubation at 37 °C (Fig 5E, orange circles and text). We conclude host-cell temperature to play a key determinant in the differential regulation of MPXV transcription at both physiological skin (33 °C) and febrile (38.5 or 40 °C) temperatures relative to core body temperature (37 °C). Collectively (Figs 2 to 5), these data identify the optimal temperature for MPXV Clade IIb.B1 replication to be 37°C.

**Fig 5.**
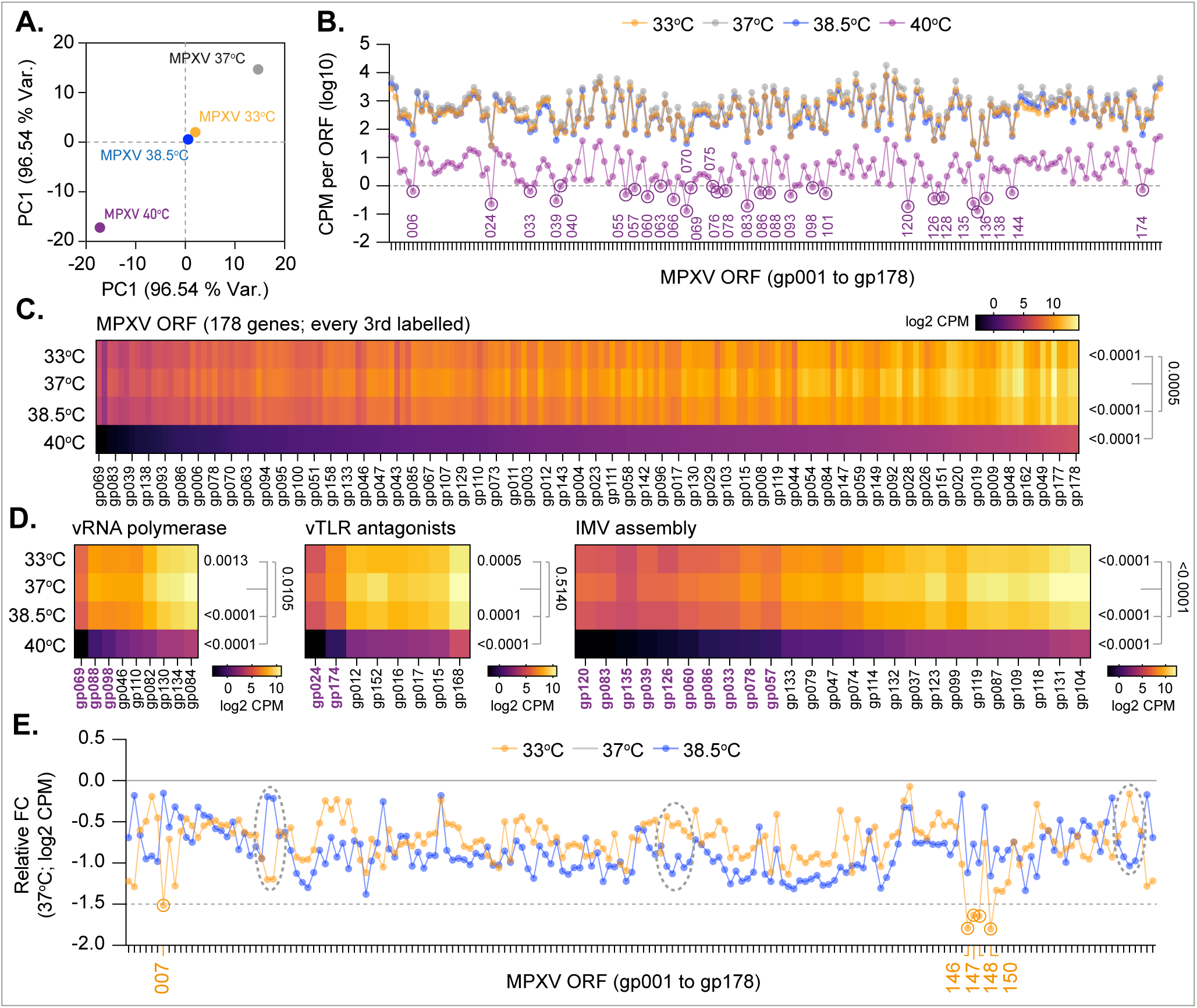
Basal host-cell temperature differentially influences the regulation of MPXV transcription. HFt cells were infected with MPXV (MOI 0.01 PFU/cell) for 1 h at 37 °C prior to incubation at 33, 37, 38.5 or 40 °C. RNA was extracted at 48 h post-infection (hpi) for RNA-Seq. Viral reads were aligned to the MPXV reference sequence ([NC_063383.1](https://www.ncbi.nlm.nih.gov/nuccore/NC_063383)) and normalised to counts per million (CPM) and used to quantify MPXV transcript expression levels per ORF. (A) Principal Component (PC) analysis of MPXV ORF transcript expression levels per incubation temperature. (B) Line graph showing CPM per ORF (log10). Purple circles and text highlight viral ORFs < 1 CPM (dotted line). (C) Expression level of MPXV ORFs (log2 CPM); every third ORF labelled. (D) Expression level of MPXV ORFs relating to the viral RNA (vRNA) polymerase complex, viral toll like receptor (vTLR) antagonists, and intracellular mature virus (IMV) assembly (as indicated). Purple text highlights viral ORFs < 1 CPM per ORF (as in B). (E) Line graph showing normalised CPM count per ORF (log2 CPM) at 33 and 38.5 °C. Values were normalised to CPM counts at 37 °C (solid grey line). Orange circles and text highlight viral ORFs ≥ 1.5 log2 FC (fold change) relative to 37 °C (dotted grey line). (A to E) *N*=3 independent biological experiments; *p*-values shown, Dunnett’s paired one-way ANOVA (bottom), paired two-tailed *t* test (top). Raw values presented in supplemental S5 data.

### Basal temperature differentially regulates the host-cell response to MPXV infection

As we had identified temperature to differentially regulate MPXV transcription (Fig 5), we next investigated the influence of temperature on the host-cell response to infection. We performed RNA-Seq analysis on mock-treated or MPXV-infected HFt cells incubated at 33, 37, 38.5 or 40 °C (MOI 0.01 PFU/cell, 48 h). Pairwise comparisons identified unique and shared clusters of DEGs (FDR < 0.05, ≥ 1.5 log2 FC) across the temperature range of analysis (Fig 6A, B). Relatively few DEGs were identified between mock-treated and infected samples incubated at 40 °C, suggesting MPXV infection at this temperature to induce a minimal host-cell response (Fig 6A, B; yellow ellipses and lines). This was surprising, as we had observed equal levels of MPXV genome entry into infected cells across the temperature range of analysis (Fig 3J). These data identify the onset of MPXV replication to be a key determinant in the host-cell response to infection. Pathway analysis identified distinct profiles of DEG enrichment dependent on incubation temperature, with 33 and 38.5 °C sharing the highest degree of similarity in their respective host-cell responses to infection (Fig 6C, grey lines). Notably, many pathways were only significantly enriched in response to infection at 37 °C (Fig 6C), even though cells incubated at 33 and 38.5 °C were observed to be productively infected and to be releasing infectious virus (Figs 2 to 4). These data demonstrate basal host-cell temperature to play a key role in the outcome and/or amplitude of host-cell response to MPXV infection. Consistent with this finding, we observed multiple cytokine-related pathways to be differentially regulated in response to MPXV infection (Fig 6C, black arrows). Analysis of these DEGs (71 genes in total) revealed distinct profiles of immune gene expression that were dependent on both infection and incubation temperature (Fig 6D to F). Importantly, this host-cell immune signature to infection occurred independently of the robust induction of the type-I IFN response (Fig 6G; Fig S4A, B). While *IFNB1* was identified to be a DEG (Fig 6G, black arrow), levels of *IFNB1* expression were extremely low (MPXV 37 °C mean CPM = 0.1285, -/+ 0.02 SD; Fig S4A, B), with little to no change observed in the overall profile of host interferon responsive genes (IRGs; Fig S4C, D). These data demonstrate Clade IIb.B1 MPXV infection to effectively suppress the induction of IFN-mediated antiviral immune defences. Notably, MPXV-infected cells incubated at 40 °C expressed an equivalent cytokine profile to that of mock treatment at this incubation temperature (Fig 6D, E [purple arrow]; Fig S4E). Thus, viral genome entry into host-cells alone is not sufficient to trigger the activation of pathogen recognition receptors (PRRs) necessary for the induction of cytokine-mediated pro-inflammatory immune responses to infection. Collectively, these data demonstrate MPXV to induce a temperature-dependent cytokine response to infection dependent on the productive onset of viral lytic replication (Fig 3F to I).

**Fig 6.**
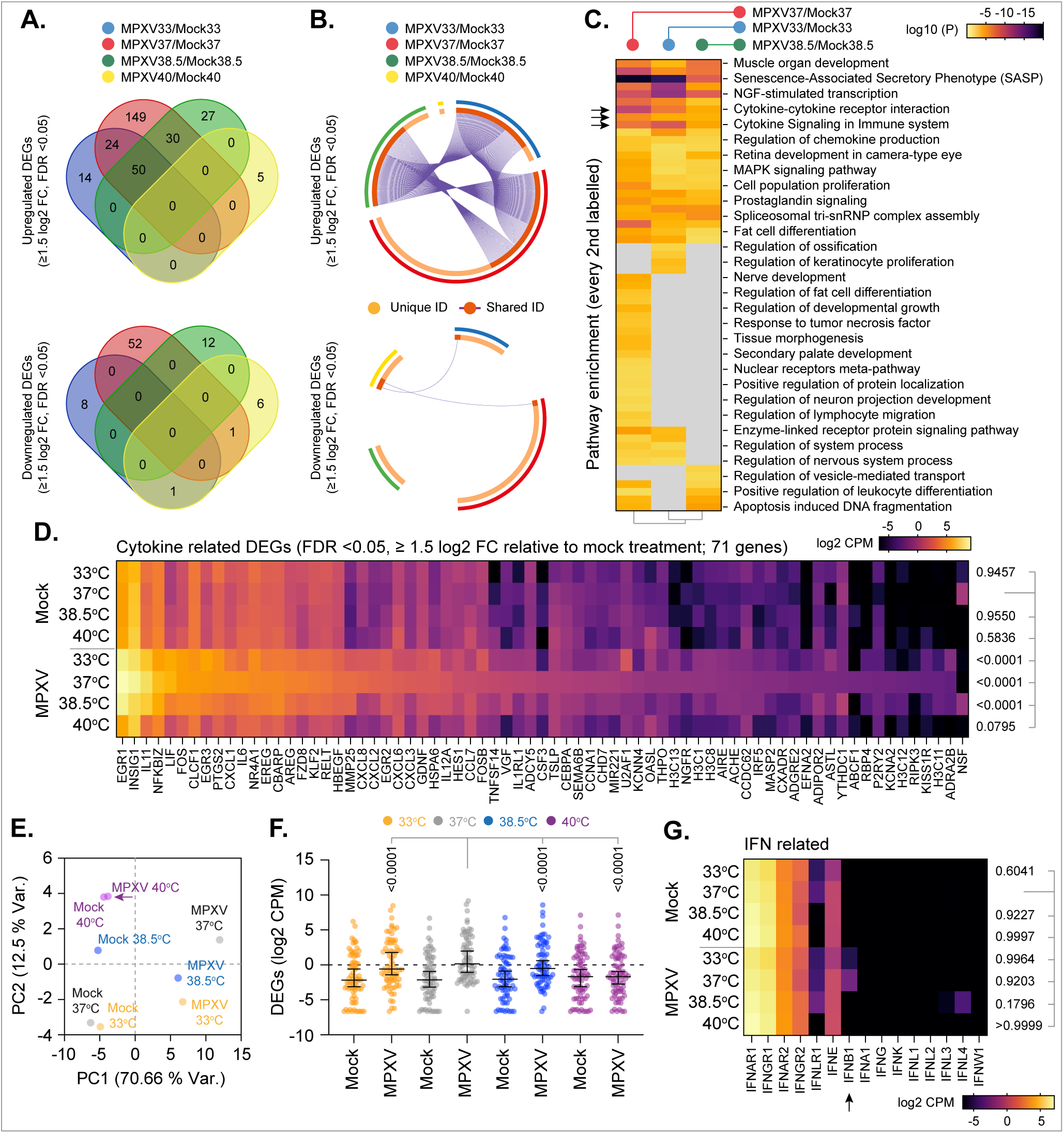
Basal temperature influences the host-cell response to MPXV infection. HFt cells were mock treated or infected with MPXV (MOI 0.01 PFU/cell) for 1 h at 37 °C prior to incubation at 33, 37, 38.5 or 40 °C. RNA was extracted at 48 h post-infection (hpi) for RNA-Seq. Host mapped reads were aligned to the human genome, normalised to counts per million (CPM), and DEGs (FDR < 0.05, ≥ 1.5 log2 FC) identified for each paired condition analysed (as indicated). (A) Venn diagram showing the number of unique or shared DEGs between each paired condition. (B) Circos plots showing the proportion of unique (light orange inner circle) or shared (dark orange inner circle + purple lines) DEGs between each paired condition. (C) Metascape pathway analysis showing significant up-regulated DEG enrichment (*p*-value < 0.05; log10 *p*-values shown) for each paired condition (as in A/B). Black arrows highlight enriched for cytokine related pathways; grey boxes, *p*-value > 0.05. (D) Expression level (log2 CPM) of cytokine related DEGs identified to be upregulated in response to MPXV infection (black arrows in C, 71 genes in total; log2 CPM). (E) Principal Component (PC) analysis of cytokine related DEGs (as in D) in mock treated and MPXV infected cells (as indicated). Purple arrow highlights MPXV 40 °C expression levels clustering with mock treated samples. (F) Expression profile of cytokine related DEGs (as in D). Black line, median; whisker, 95% CI; all data points shown. (G) Expression level (log2 CPM) of host interferon (IFN)-related receptors and cytokines in mock treated or MPXV infected cells. Black arrow highlights expression level of *IFNB1* in mock treated or MPXV infected cells. (D, F, G) *P*-values shown, Dunnett’s paired one-way ANOVA. (A to G) *N*=3 independent biological experiments. Raw values presented in supplemental S6 data.

Investigating further, we compared the host-cell transcriptome profiles of MPXV infected cells incubated at 37 and 40 °C. Out of the 935 DEGs (FDR <0.05, ≥ 1.5 log2 FC) identified (Fig 7A), pathway analysis identified cell cycle (Fig 7B, grey arrow and circle) and immune system (Fig 7B, coloured arrows and circles) related pathways to be predominantly downregulated at 40 °C relative to incubation at 37 °C (Fig 7B). Analysis of immune system related DEGs (61 genes in total) identified signalling by interleukin (IL; 20 DEGs) and adaptive immune system (21 DEGs) pathways to account for the majority of DEGs identified between these two infection conditions (Fig 7C, D). In contrast to IFN signalling (Fig S4A to D), analysis of the IL signalling pathway (R-HAS-449147; 459 genes in total) identified this pathway to be significantly upregulated during MPXV infection in a temperature-dependent manner (Fig 7D green dots; Fig S4F). Notably, multiple cytokines and chemokines implicated in IL signalling were observed to vary in their relative profiles of expression between 33 and 38.5 °C (Fig 7E; e.g., IL12A [*P* = 0.0026], CXCL2 [*P* = 0.0013]). Importantly, MPXV incubation at 40 °C failed to upregulate these cytokine-related genes above baseline levels (Fig 7D, E; Fig S4E, F). Together, these data demonstrate MPXV to induce a robust IL response to infection in a temperature-dependent manner.

**Fig 7.**
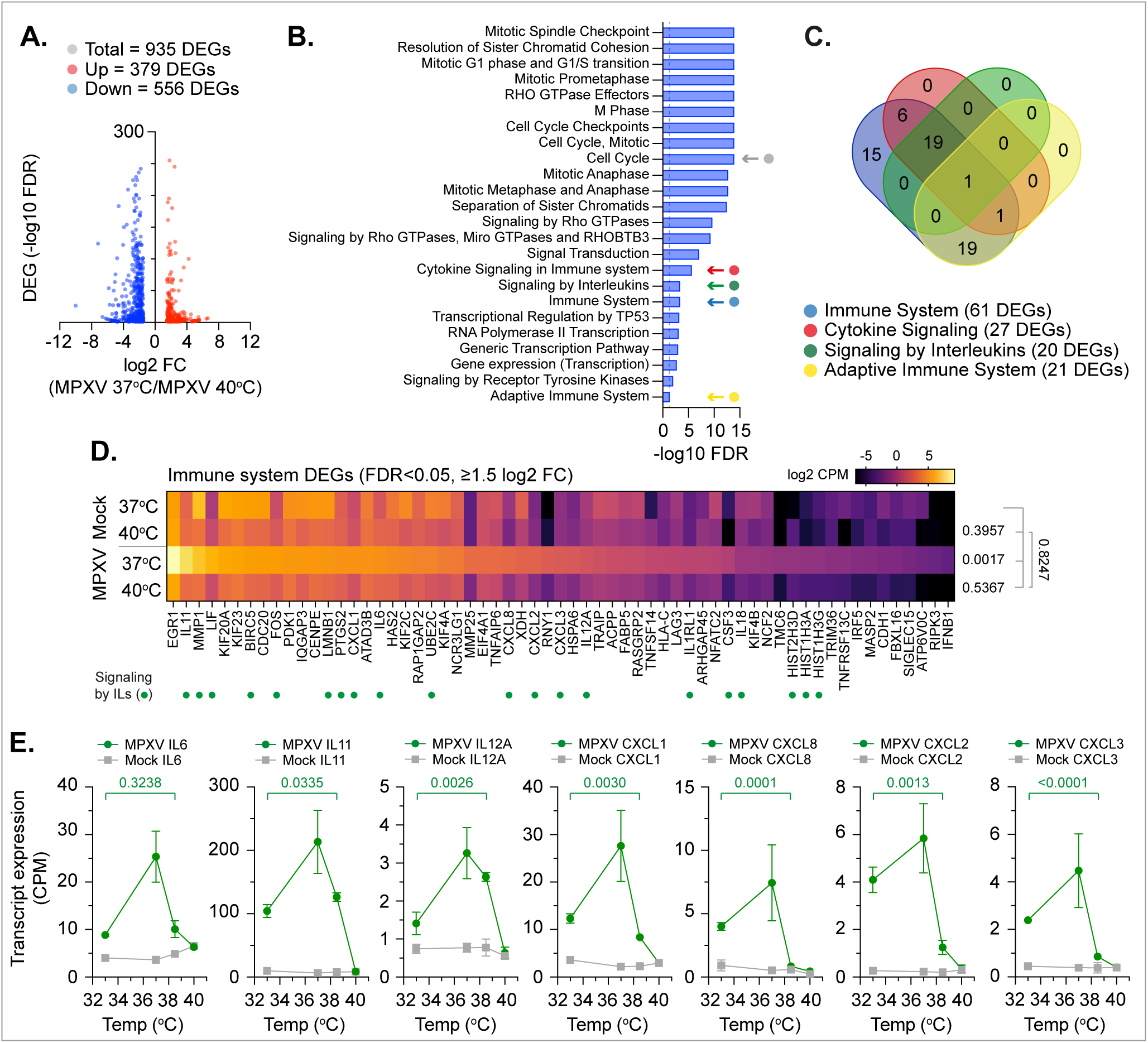
Basal host-cell temperature influences the interleukin response to MPXV infection. HFt cells were mock treated or infected with MPXV (MOI 0.01 PFU/cell) for 1 h at 37 °C prior to incubation at 33, 37, 38.5 or 40 °C. RNA was extracted at 48 h post-infection (hpi) for RNA-Seq. Host mapped reads were aligned to the human genome, normalised to counts per million (CPM), and DEGs (FDR < 0.05, ≥ 1.5 log2 FC) identified. (A) Scatter plot showing DEGs identified between MPXV infected cells incubated at 37 and 40 °C; up-regulated, red circles; down-regulated, blue circles. (B) Reactome pathway analysis of down-regulated mapped DEGs (blue circles in A). Top 25 down-regulated pathways (FDR < 0.05) shown (blue bars; plotted as -log10 FDR). Dotted line, threshold of significance (-log10 FDR of 0.05). Pathways relating to cell cycle (grey arrow and circle) and immune system regulation (coloured arrows and circles) are highlighted. (C) Venn diagram showing the number of unique or shared DEGs identified in immune system pathways (coloured arrows and circles identified in B). (D) Expression level (log2 CPM) of immune system DEGs (identified in B; 61 genes in total); *p*-values shown, Dunnett’s unpaired one-way ANOVA (top), unpaired two-tailed *t* test (bottom). DEGs associated with Signalling by Interleukins (ILs) highlighted (green circles). (E) Expression profile (CPM) of selected interleukins (IL6, IL11, and IL12A) and chemokines (CXCL1, CXCL6, CXCL2, CXCL3) in mock treated (grey lines) and MPXV infected (green lines) cells across the temperature range of analysis; *p*-values shown, unpaired two-tailed *t* test. Raw values presented in supplemental S7 Data.

### Basal temperature differentially regulates the host-cell IFN response to MPXV infection

As the IFN response is known to play a key role in limiting OPXV pathogenicity and immune clearance *in vivo*^44,45^, we next investigated whether basal host-cell temperature influenced the IFN-mediated host-cell restriction of MPXV. HFt cells were stimulated with IFN-β and incubated at 33, 37, 38.5 or 40 °C for 16 h prior to MPXV infection (MOI 0.001 PFU/cell) and continued incubation in the presence of IFN for 48 h at their respective temperatures. Quantitation of plaque counts demonstrated MPXV restriction to occur in an IFN-β dose- and temperature-dependent manner, with heightened levels of restriction observed at 38.5 °C relative to incubation at 33 or 37 °C (Fig 8A to D). Consistent with our previous analysis (Figs 2 to 4), MPXV plaque formation was abrogated at 40 °C irrespective of IFN stimulation (Fig 8A, B grey dotted lines and circles). IFN pre-treatment and infection with VACV demonstrated an analogous dose- and temperature-dependent trend in VACV restriction that abrogated plaque formation at 40 °C in the presence of 100 IU/ml of IFN-β (Fig S5A to D). As temperature elevation alone led to suppressed levels of MPXV transcription independently of the robust induction of the type-I IFN response (Figs 5, 6, S4A to D), we next investigated the synergistic nature of temperature elevation and IFN co-stimulation on the outcome of MPXV infection. Naïve HFt cells were infected with MPXV (MOI 0.01 PFU/cell) for 1 h at 37 °C prior to stimulation with IFN-β and incubation at 33, 37, 38.5 or 40 °C. Infected and treated cell monolayers were fixed at 24 hpi and the number of MPXV virion antigen positive cells quantified by immunostaining. Consistent with our IHC analysis of infected tissues (Fig 2), incubation temperature dramatically reduced the number of MPXV virion antigen positive cells detected at 24 hpi (Fig 8E, F; 0 IU/ml IFN-β, 37°C > 33 ≈ 38.5 °C > 40 °C). Notably, a substantial number of MPXV antigen positive cells could still be detected at 40 °C, indicating that a proportion of MPXV genomes were able to circumvent complete thermal restriction (Fig 8E, F; 0 IU/ml IFN). Co-stimulation with IFN-β led to a dose- and temperature-dependent reduction in the number of antigen positive cells (Fig 8E, F plus IFN). These data identify temperature elevation to cooperate and synergise with the type-I IFN response to mediate the host-cell restriction of MPXV. As neither temperature elevation nor IFN stimulation was sufficient to eliminate MPXV gene expression (Fig 8E, F), we hypothesised MPXV replication may recover following temperature downshift to 37 °C and/or withdrawal of IFN as an immune stimulus. Downshift of infected cells from 40 (no IFN treatment) to 37 °C led to a complete recovery in MPXV plaque formation to equivalent levels observed at 37 °C (Fig 8G). These data demonstrate MPXV thermal restriction to only stall the progress of infection, as opposed to eliminate or induce the establishment of viral genome quiescence. Surprisingly, IFN washout from infected and IFN-treated cells incubated at 37 °C also led to an equivalent recovery (Fig 8H). Analogous to temperature restriction, these data demonstrate the type-I IFN response to suppress, but not eliminate, MPXV infection. Importantly, IFN washout and thermal downshift of MPXV-infected and IFN-stimulated cells from 40 to 37 °C led to significantly lower recovery in the re-establishment of MPXV infection in an IFN dose-dependent manner (Fig 8J). Equivalent trends in the re-establishment of infection were also observed following IFN washout and temperature downshift of VACV infected cells (Fig S5G to J). We conclude temperature elevation to synergise with and enhance the type-I IFN-mediated restriction of MPXV to limit the re-establishment of infection upon homeostatic recovery from these two independent pro-inflammatory immune responses known to be activated in response to infection *in vivo*^32,46^.

**Fig 8.**
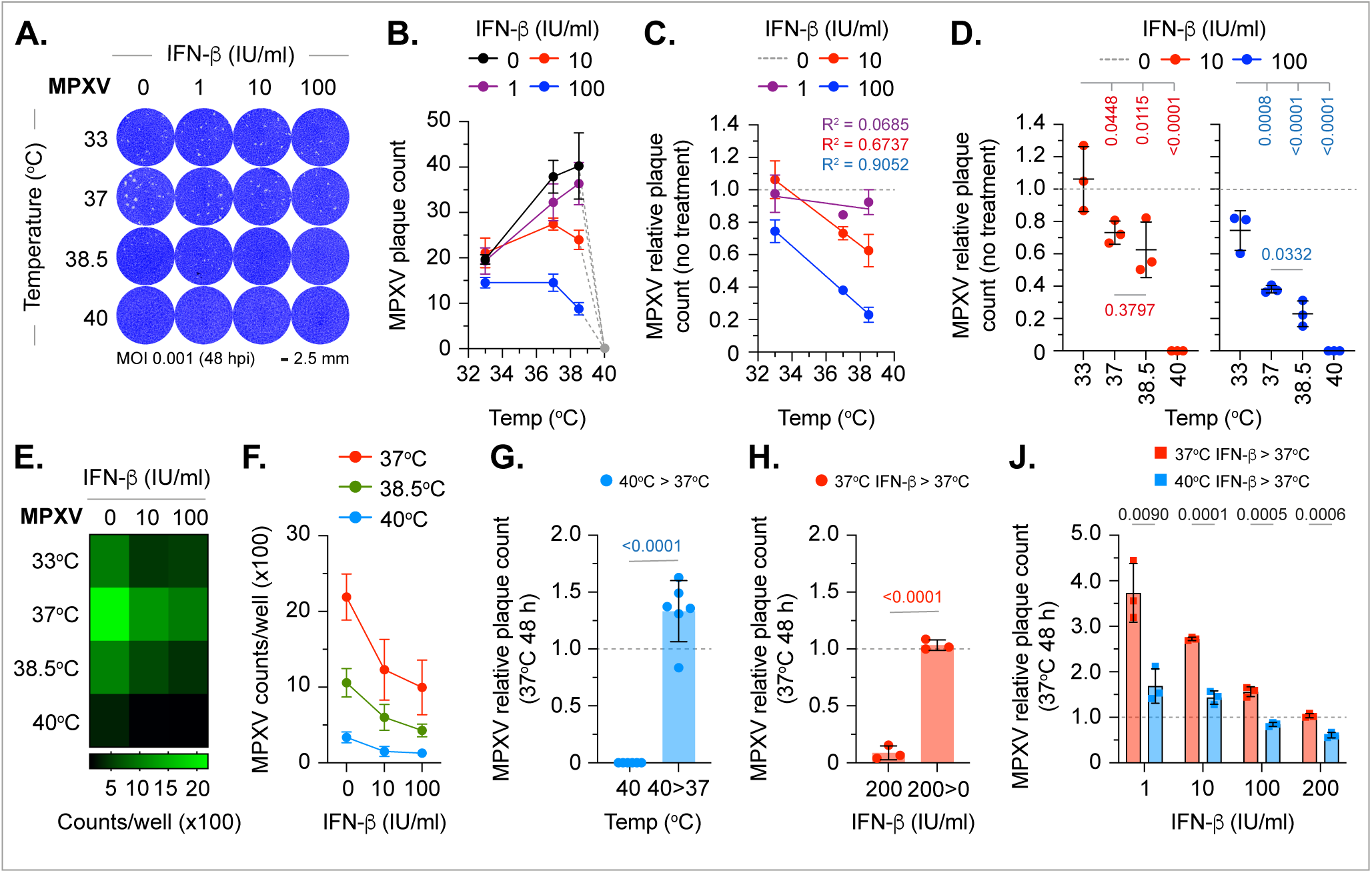
Temperature elevation enhances the type-I IFN-mediated host-cell restriction of MPXV. (A to D) HFt cells were pre-treated with IFN-Π (0 to 100 IU/ml, as indicated) for 16 h at 33, 37, 38.5 or 40 °C prior to MPXV (MOI 0.001 PFU/cell) infection (1 h at 37 °C) and continued incubation at their respective incubation temperatures in the presence of IFN. (A) Representative images of MPXV infected cell monolayers pre-treated with IFN and stained with Coomassie Brilliant blue at 48 h post-infection (hpi). (B) Quantitation of MPXV plaque counts at 48 hpi (as shown in A). Means and SD shown. Grey dotted lines and circles highlight absence of plaque formation at 40 °C. (C/D) Relative MPXV plaque counts. Values were normalised to no IFN treatment (dotted grey line) per incubation temperature. (C) Means and SD shown; coloured lines and text, linear regression and corresponding R^2^ values. (D) As in C, all data points shown; black line, mean; whisker, SD; *p*-values shown, Dunnett’s unpaired one-way ANOVA. (E to J) Naïve HFt cells were infected with MPXV (MOI 0.01 PFU/cell) for 1 h at 37 °C prior to overlay with media containing IFN-Π (0 to 200 IU/ml, as indicated) and incubation at 33, 37, 38.5 or 40 °C. (E/F) Cells were fixed at 24 h and stained for MPXV virion protein expression and the number of antigen positive cells quantified by indirect immunofluorescence. (E) Mean MPXV positive cell counts (x100) per infected cell monolayer. (F) As in E, mean and SD shown. (G to J) Infected or infected and IFN treated monolayers incubated at 37 or 40 °C were either fixed at 48 h or washed twice and overlayed with fresh media (IFN washout) and incubation at 37 °C for an additional 48 h prior to fixation at Coomassie Brilliant blue staining. (G) Quantitation of MPXV plaque counts in cell monolayers incubated at 40 °C or temperature downshifted from 40 to 37 °C (40>37) with continued incubation. (H) Quantitation of MPXV plaque counts in cell monolayers incubated at 37 °C in the presence of IFN or following IFN washout and continued incubation. All data points shown. (J) Quantitation of MPXV plaque counts in cell monolayers incubated at 37 or 40 °C in the presence of IFN (1 to 200 IU/ml, as indicated) with IFN washout at 48 h and continued incubation at 37 °C (no IFN). All data points shown. (G to J) Values were normalised to plaque counts determined at 37 °C (no IFN treatment at 48 h). Mean and SD shown; *p*-values shown, unpaired two-tailed *t* test. Raw values presented in supplemental S8 Data.

## Discussion

The emergence of novel MPXV Clade IIb variant strains that support heightened levels of human-to-human transmission is a global health concern and highlights the need for a better understanding of MPXV and its interaction with the host immune system. The pro-inflammatory fever response is a well-established and common prodrome symptom of MPXV infection^11,14-17^, but the precise thermoregulatory impact on the outcome of infection has remained poorly defined. Here, we isolated a clinical Clade IIb.B.1 MPXV strain (MPXV CVR_S1) from a hospitalised patient admitted with fever (38.5 °C) to determine the influence of temperature elevation on the outcome of MPXV infection.

Previous studies (circa 1960s) have shown OPXVs to have divergent ceiling temperatures of pox formation in the chorioallantoic membrane of infected embryonic chicken eggs, ranging from 38 °C to ≥ 40.5 °C (VARV minor to VACV, respectively)^33,34^. Here we show the clinical febrile temperature range of 38.5 to 40 °C, representative of low- to moderate-grade fever in humans^11,14-17,21^, to restrict the replication of a circulating MPXV Clade IIb strain. This temperature-dependent restriction was observed across a variety of infection models (Figs 2 to 4), including pseudostratified skin epithelium, and is consistent with the reported ceiling temperature of MPXV pox formation in embryonic chicken eggs of 39.5 °C^33^. Collectively, these data demonstrate that febrile temperatures associated with mpox disease in humans are likely to be a key determinant in the outcome of infection. Importantly, we show that the thermal restriction of MPXV to occur independently of a block to virus entry (Fig 3J) and to be reversible upon temperature downshift without impairment to viral replication fitness (Fig 8G). These findings are consistent with the re-establishment of VARV replication in embryonic chicken eggs upon thermal downshift from its inhibitory ceiling temperature of 38.5 °C^34^. As OPXVs with alternate ceiling temperatures have been shown to complement quiescent infections leading to hybrid pox formation^34,47^, variance in MPXV Clade and strain ceiling temperatures might contribute to genetic reassortment during mixed co-infection and/or changes in viral pathogenesis. Thus, the thermoregulatory mechanisms surrounding MPXV genome stability, longevity, and retention of replication fitness at febrile temperatures warrants further investigation. Importantly, we demonstrate lower physiological temperatures associated with skin tropism to also impact on MPXV replication. We observed a significant reduction in MPXV intraepithelial propagation, plaque formation, and virus yield following incubation at 33 °C (Figs 2 to 4). We posit the fever response to both positively and negatively influence the outcome of OPXV infection dependent on the baseline temperature of the tissue at the point of infection. For example, a general two to four °C rise in skin temperature from 33 to 37 °C would be expected to promote MPXV replication, whereas a similar rise in tissue temperature from 37 °C we show to be progressively inhibitory (Figs 2 to 4). Thus, inherent differences in species body/tissue temperature and/or capacity to mount a febrile immune response could impact on the zoonotic potential of MPXV to be maintained within an animal reservoir or to undergo inter-species host transmission. Indeed, available evidence suggests small rodent models to have variable febrile responses to OPXV infection, potentially in an MPXV Clade dependent manner^22,30-32^. Thus, the influence of tissue temperature on MPXV species tropism and onward transmission also warrants additional investigation.

Our transcriptomic analysis demonstrates host-cell temperature to have a significant impact on the differential regulation of viral transcription and associated host-cell responses to MPXV infection, both at physiological (33 °C) and febrile (38.5 and 40 °C) temperatures relative to core body temperature (37 °C; Figs 5 to 7). MPXV incubation at 40 °C led to a substantial decrease in viral transcription (≈ 100-fold reduction per ORF; Fig 5B, C). However, the overall profile of transcription remained proportionate to that observed at 37 °C (Fig 5B), indicative of a genome-wide suppression in transcription, as opposed to a specific block in the expression of any individual viral ORF. Notably, we identify multiple viral ORFs that encode proteins associated with virion maturation to show substantially decreased transcript levels (< 1 CPM per ORF; Fig 5B, D). Consistent with this observation, significantly lower levels of virion antigen expression were also observed within MPXV-infected tissues or cells incubated at 40 °C (Figs 2, 8E). Thus, temperature elevation is likely to cause a bottleneck in the expression of critical viral gene products required for virion maturation that suppress the onward propagation and spread of MPXV. Notably, differential patterns of MPXV transcription and replication fitness were also observed at lower temperatures (Figs 2, 3, 4, 5E). Incubation at 33 °C led to a 50 % decrease in viral plaque formation, whereas incubation at 38.5 °C resulted in a 50 % decrease in viral plaque size relative to incubation at 37 °C (Figs 3G, H, 4C). Together, these data highlight the importance of basal host-cell temperature to the outcome of MPXV infection. We posit that the differential patterns of viral transcription observed between 33 and 38.5 °C to account for the phenotypic differences identified between these two incubation temperatures. Of the five viral ORFs identified to be significantly downregulated at 33 °C (Fig 5E, orange circles), experimental evidence has shown the VACV orthologues of gp148 (VACV A43R), gp007 (VACV D6L), and gp146 (VACV A41L) to impact on viral replication fitness and lesion formation^48-51^. Thus, the differential patterns of viral transcription observed at both physiological and febrile temperatures are likely to affect multiple viral and cellular processes that govern the overall outcome of infection.

Our host-cell transcriptome analysis of mock-treated or MPXV-infected cells incubated at 40 °C identified relatively few DEGs at this incubation temperature (13 in total; Fig 6A, B). This minimal host-cell response correlates with the limited levels of MPXV transcription and replication observed at 40 °C (Figs 2 to 5). Thus, viral genome entry into host-cells alone is not sufficient to activate PRRs sufficient to induce a robust pro-inflammatory immune response to infection (Figs 6, S4). In contrast, we identify a significant induction of pro-inflammatory interleukin gene expression at both physiological skin, core, and febrile temperatures (33, 37 and 38.5 °C, respectively) permissive to MPXV replication. These differential patterns of interleukin and chemokine expression correlate well with the levels of MPXV replication observed at these incubation temperatures. For example, IL-6 induction (a key pyrogen in the regulation of the fever response^52^) demonstrated peak induction at 37 °C, with lower levels of induction observed at 33 or 38.5 °C (Fig 7E), consistent with decreased levels of replication observed at these incubation temperatures (Figs 2, 3). Thus, we identify basal host-cell temperature to be a key determinant in the amplitude of cytokine response to MPXV infection. These data are consistent with cytokine profiling experiments of serum samples derived from experimentally infected NHPs or hospitalised patients that have shown pro-inflammatory interleukin expression levels to correlate with the kinetics of MPXV replication and disease severity, respectively^32,46^. While MPXV encodes immune antagonists to IL-1 (gp014, gp152, gp167), IL-18 (gp007), TNF-α (gp002, gp178), IFN-α/μ (gp169), and IFN-γ (gp163), among others^44^, it does not encode a direct IL-6 antagonist. This raises the possibility that specific profiles of pro-inflammatory cytokine or chemokine expression may be beneficial to MPXV replication prior to temperature elevation, potentially through the recruitment and infection of bystander and/or circulating immune cells to promote virus dissemination^44^.

The type-I IFN response is a critical component of the host’s antiviral immune defence, which acts to restrict viral propagation and to prime adaptive immune responses to clear MPXV infection^44,45,53-55^. Due to their large coding capacity, OPXVs encode multiple immune antagonists to block the induction of the type-I IFN response at several independent stages of infection^44,45,56,57^. Correspondingly, we demonstrate a clinical and circulating MPXV IIb.B1 strain to effectively suppress the induction of the type-I IFN response (Figs 6, S4). Importantly, the activation of PRRs that regulate the pro-inflammatory IFN response can be triggered through multiple mechanisms, including the detection of damage-associated molecular patterns (DAMPs) in addition to pathogen associate molecular patterns (PAMPs). We posit the IFN signature observed in response to MPXV infection *in vivo* is likely a consequence of DAMP activation due to the high levels of replication observed in multiple cell-types and organs upon viral dissemination^32,44,46^. We demonstrate a circulating clinical MPXV IIb.B.1 strain to be sensitive to exogenous IFN-β stimulation in a dose- and temperature-dependent manner (Fig 8). These data are consistent with previous reports that have shown Clade I MPXV strains to be sensitive to exogenous IFN stimulation at 37 °C^53,55^. Importantly, we demonstrate this IFN-mediated restriction to be reversible and not to effectively eliminate infection, with significantly lower levels of MPXV recovery observed at febrile temperatures associated with clinical infection (Fig 8H, J). Thus, we identify functional cooperativity and synergism between pro-inflammatory immune responses (fever plus IFN) that mediate the accumulative host-cell restriction of MPXV. Such accumulative effects are likely to be a consequence of suppressed levels of viral transcription at febrile temperatures restricting the ability of MPXV to express immune antagonists that directly counteract the effects of IFN (e.g., the viral IFN-α/μ receptor antagonist gp169)^56^. We posit that such synergism between pro-inflammatory immune responses will likely play a significant role *in vivo* to limit MPXV propagation and to prime adaptive immune responses required for immune clearance^54^.

In conclusion, our study provides critical insights into the importance of temperature in the regulation and outcome of MPXV infection. We identify host-cell temperature at both physiological and febrile temperatures to be a key determinant in the regulation of viral transcription, associated amplitude of cytokine response to infection, and ancillary interactions with the type-I IFN response required to inhibit MPXV replication. Our data shed light on the complex interplay and functional synergy between host pro-inflammatory immune systems and their net combinatorial affect on the outcome of OPXV infection, findings pertinent to the immune regulation of many clinical pathogens that induce a fever response.

## Materials and Methods

### Cells and cell viability assays

Vero E6 (a gift from Michelle Bouloy, Institute Pasteur, France), CV-1 (European Collection of Authenticated Cell Cultures (EACC), 87032605), HFt^58^, RPE-1 (ATCC, CRL-4000), HaCaT (Addexbio, T0020001), and J2 3T3 (a gift from Sheila Graham, MRC-University of Glasgow Centre for Virus Research) cells were grown in Dulbecco’s Modified Eagle Medium (DMEM; ThermoFisher, 10566016) supplemented with 10 % fetal bovine serum (FBS; ThermoFisher 10270106) and 1 % penicillin streptomycin (PS; ThermoFisher, 15070063). NOK cells (a gift from Karl Munger, Tufts University School of Medicine, USA)^59^ were grown in Keratinocyte-SFM medium with L-glutamine, EGF, and BPE (ThermoFisher, 17005075). All cells were grown and maintained at 37 °C in 5 % CO_2_ unless otherwise stated. Cell viability was measured using MTS reagent (Abcam, ab197010), as per the manufacturer’s guidelines, and absorbance measurements taken at OD=490 nm using a PHERAstar plate reader (BMG LABTECH). As a positive control for cell death, puromycin (Sigma-Aldrich, P8833) was added to cell culture media at a final concentration of 1.0 mg/ml and incubated for 24 h prior to the addition of MTS.

### Organotypic raft culture

Type I collagen was extracted from rat tails (supplied by the University of Glasgow, Veterinary Research Facility). Ethanol sterilised tails were incubated in 0.5 M acetic acid at 4 °C for 48 hours on a magnetic stirrer. Material was clarified by centrifugation at 10,000 x g for 30 minutes at 4 °C. Supernatant was subjected to collagen extraction by NaCl precipitation (final concentration 10 % weight/volume) for 1 h at 4 °C and collected by centrifugation at 10,000 x g for 30 minutes at 4 °C. The pellet was resuspended overnight in 0.25 M acetic acid and dialysis for three days (two changes per day) in pre-cooled acetic acid (final concentration 17.4 mM) at 4 °C. Sterile collagen was isolated by centrifugation 20,000 x g for 2 h at 4 °C. Organotypic raft cultures were prepared as previously described^60^. Briefly, collagen matrices with J2 3T3 fibroblasts were prepared in 12-well plates and allowed to contract for approximately 7 days at 37 °C. Keratinocytes (HaCaT cells) were seeded on top of contracted collagen matrices at a density of 2.5×10^5^ cells/matrix in 24-well plates and incubated at 37 °C overnight prior to being transferred into Costar Transwells (Corning Costar, 3401) and maintained at 37°C under air liquid interface (ALI) for 12 to 14 days. Tissues were maintained in E-medium (3:1 ratio of DMEM:F12 (Thermofisher, 21765029) medium supplemented with 10 % FBS and 1 % PS, 2 mM L-glutamine (ThermoFisher, 25030081), 180 μM adenine (Sigma-Aldrich, A2786), 5 μg/ml transferrin (Sigma-Aldrich, T1147), 5 μg/ml insulin (Sigma-Aldrich, I6634), 0.4 mg/ml hydrocortisone (Sigma-Aldrich, H0888), 0.1 nM cholera toxin (Sigma-Aldrich, C8052), and 0.2 ng/ml epidermal growth factor (EGF; Sigma-Aldrich, E9644). Basal-chamber E medium was changed every 48 h. Pseudostratified skin tissues were infected with 10^2^ or 10^3^ pfu/tissue (as indicated) in 50 µl inoculum for 1 h at 37 °C prior to incubation at the indicated temperatures for 72 h.

### Immunohistochemistry

Mock treated or infected pseudostratified skin epithelia was fixed in 8% (v/v) formaldehyde overnight, washed in PBS, and processed by the Diagnostic Services, School of Biodiversity, One Health and Veterinary Medicine, University of Glasgow. Following paraffin embedding, 3 µm sections were cut and placed in a 37 °C oven overnight. Samples were dewaxed in HistoClear (National Diagnostics, HS-20) and rehydrated through 3x sequential washes in ethanol (final concentration 70 %). Tissue sections were subject to antigen retrieval using TET buffer (10 mM Tris base pH 9, 1 mM EDTA, 0.05% Tween-20) in a MenaPath access retrieval unit at 125 °C and 15 psi for 100 sec, before cooling to room temp in H_2_O and transfer to TNET buffer (10 mM Tris pH 7.5, 100 mM NaCl, 1 mM EDTA, 0.05 % Tween-20). Tissue sections were stained with a polyclonal antibody raised against whole VACV (Invitrogen, PA1-7258; 1/2000 dilution) for 30 mins, washed in TNET buffer, prior to secondary staining (Dako Envision polymer anti-rabbit, K4003) for 30 mins and washing in TNET buffer. Visualisation was achieved using two 5 min applications of DAB+ (Dako, K3468). Tissue sections were counterstained using Gills haematoxylin dehydrated, prior to clearing in Histoclear, and mounting with a coverslip. Tissue sections were imaged using an Aperio VERSA Digital Pathology Scanner (Leica Biosystem). Scanned tissue sections were quantified using the Halo multiplex IHC module image analysis software (indica labs). The algorithm was adapted for nuclei identification and arbitrary thresholds applied to detect weak, moderate, and strong MPXV antigen positive cells over background levels of staining. Algorithm optimisation was performed using the real-time Tuning function. All sections were analysed using the same algorithm and subjected to random checks to confirm the precision of tissue annotation.

### Viruses

Clinical monkeypox virus (MPXV) CVR_S1 was isolated from an ISARIC4C patient with ethical consent (Ethics approval for the ISARIC CCP-UK study was given by the South Central–Oxford C Research Ethics Committee in England (13/SC/0149), the Scotland A Research Ethics Committee (20/SS/0028) and the WHO Ethics Review Committee (RPC571 and RPC572). The patient presented with fever (38.5 °C), cough, myalgia, and skin rash. The patient was haemodynamically stable and not requiring supplemental oxygen. Swabs were collected from multiple skin lesions into virus transport medium (VTM). The VTM was mixed in a 1:4 ratio with DMEM supplemented with 10 % FBS, 1 % PS and 250 ng/ml Amphotericin B (ThermoFisher, 15290018) and centrifuged at 4000 rpm for 10 mins. Vero E6 (sub-clone MESO; a gift from Meredith Stewart, MRC - University of Glasgow Centre for Virus Research) cells were inoculated with 500 μl of VTM/media for 1 h. Infected monolayers were washed three times in PBS prior to overlay with cell culture media and incubation at 37 °C in the presence of 5 % CO_2_. Media containing infectious cell-released virus (CRV; primary amplification P1 stock) was harvested at 72 hours post-infection (hpi) and clarified at 4000 rpm for 10 minutes. P1 virus was subjected to Illumina metagenomic sequencing to evaluate purity and sequence homology to pre-culture clinical material (CVR_MPXV1a; accession number ON808413) to confirm sequence identity and source of the propagated isolate. All experiments were performed in a Biosafety level 3-laboratory at the MRC-University of Glasgow Centre for Virus Research (SAPO/223/2017/1a). Vaccinia Virus (VACV A5L-EGFP, a gift from Geoffrey Smith, University of Oxford, England)^61^ was propagated and titrated on Vero E6 (MESO) cells, as previously described^61,62^. Wild-type herpes simplex virus 1 (HSV-1, strain 17syn+; a gift from Roger Everett, MRC – University of Glasgow Centre for Virus Research) was propagated in RPE cells and titred on U2OS cells, as previously described^63^.

### Viral genome sequencing

20 ng of nucleic acid was enriched for viral genomic sequences using NEBNext Microbiome DNA Enrichment Kit (NEB, E2612) to reduce host genomic DNA contamination. Purified DNA was sheared into 350-base pair fragments by sonication using the Covaris Sonicator LE220. Library preparation was conducted with the KAPA Hyper kit (Roche, 07962347001) with end repair and universal adapter ligation. Unique Dual Index Primer pairs (NEBNext Multiplex Oligos for Illumina) were used to index the samples, followed by 10 cycles of PCR amplification. Libraries were sequenced on the Illumina NextSeq500 platform with paired ends for 2 x 150-base pair reads using a Mid-output cartridge Kit v2.5 (Illumina, 20024905). bcl2fastq software was used to demultiplex and compress bcl files into fastq (gz) files. Short and low-quality sequence reads (length < 75 nucleotides, Phred score < 30) were filtered from the fastq files using Trim Galore (version 0.6.6). BWA-MEM (version 0.7.17- r1188) was used to map filtered reads to the reference sequence. Ivar (version 1.3.1) program generated consensus sequences (minimum depth 10 reads and consensus frequency threshold 0.6) from the aligned BAM files. Consensus sequences were manually checked, and repeat regions were curated based on majority read consensus. Multiple sequence alignment was performed using MAFFT (v7.475)^64^ and aligned to the MPXV Clade IIa reference sequence ([NC_063383.1](https://www.ncbi.nlm.nih.gov/nuccore/NC_063383)). Variations between the sequences were plotted using the [Snipit program](https://github.com/aineniamh/snipit) (v.1.1.2). APOBEC3-associated mutations (G=>A and C=>T) were calculated from pairwise sequence alignment using a BASH script.

### Plaque and virus yield assays

Cells were seeded onto plates and allowed to become confluent before infection at the specified multiplicity of infection (MOI) for 1 h at 37 °C prior to overlay and incubation at the indicated temperatures. For infectious cell-released virus (CRV) quantitation, harvested supernatants were serially diluted in cell culture media and used as an inoculum to infect Vero E6 (MESO) cells to obtain a linear range of countable plaques. Infected monolayers were incubated at 37 °C for 24 to 72 h (dependent on the virus and experiment), fixed with 4 (VACV) or 8 (MPXV) % (v/v) formaldehyde (Sigma-Aldrich, F8775), washed in PBS, and stained with 0.1 % Coomassie Brilliant Blue solution (Sigma-Aldrich, B0149) in 45 % methanol and 10 % acetic acid for up to 30 minutes. Plates were washed in water and dried overnight before plaque counting under a plate microscope. For quantitation of plaque diameters, infected cell monolayers stained with Coomassie Brilliant Blue were imaged using a Celigo Imaging Cytometer (Nexcelom Bioscience, UK) and plaque measurements performed using Zen Blue software (Zeiss).

### Temperature and Interferon (IFN) inhibition assays

Cells were infected at 37 °C for 1 h at the indicated MOI prior to overlay and continued incubation at the desired temperature (33, 37, 38.5, or 40 °C) in 5 % CO_2_ for the indicated times prior to fixation and Coomassie Brilliant Blue or indirect immunofluorescence staining. For IFN inhibition assays, cell monolayers were either pre-treated with IFN-β (R&D systems, 8499-IF-010; 0.1 to 200 IU/ml, as indicated) for 16 h at the indicated temperatures prior to infection or infected for 1 h at 37 °C prior to the addition of IFN-β to the overlay and incubation at the desired temperature (as indicated). For temperature downshift and IFN washout experiments, duplicate infected plates were overlayed with media containing IFN-β (0 to 200 IU/ml; as indicated) for 24 or 48 h (VACV or MPXV, respectively) prior to either fixation or IFN washout (3x 1 ml cell culture media) and continued incubation at 37 °C for 24 or 48 h prior to fixation and Coomassie staining.

### Indirect immunofluorescence

Mock treated, infected, or infected and IFN treated cells were fixed with 4 (VACV) or 8 (MPXV) % (v/v) formaldehyde, washed in PBS, and permeabilised with 0.5 % Triton-X-100 (Sigma-Aldrich, T-9284) prior to blocking in filter sterilised PBS containing 2 % FBS (PBSf). Cells were stained with a polyclonal antibody raised against whole VACV (Invitrogen, PA1-7258; 1/2000 dilution) and secondary staining using a donkey anti-rabbit AlexaFluor 555 (Invitrogen, A31572). Cell nuclei were stained with DAPI (Sigma-Aldrich, D9542). PBSf was used for all antibody incubations (1 h at RT) and cell washing throughout. Cell monolayers were imaged and analysed using a Celigo Imaging Cytometer and companion software (Nexcelom Bioscience, UK).

### Quantitative PCR (qPCR)

PCR was carried out using NEB Luna Universal Probe One-Step RT-qPCR Kit (New England Biolabs, E3006) or Taqman Fast Universal PCR master mix (Applied Biosystems, 4352042), according to the manufactures’ instructions. VACV was detected using primers and probe directed to ORF E9L^65^; E9L forward primer: 5’-CGGCTAAGAGTTGCACATCCA-3’, E9L reverse primer: 5’-CTCTGCTCCATTTAGTACCGATTCT-3’ and E9L Probe 5’-[6FAM]AGGACGTAGAATGATCTTGTA[BHQ1]-3’. MPXV was detected using the primers and probe directed against the ORF G2R^66^; G2R FWD: 5’-GGAAAATGTAAAGACAACGAATACAG-3’, G2R REV 5’-GCTATCACATAATCTGGAAGCGTA-3’ and G2R Probe 5’-[6FAM]AAGCCGTAATCTATGTTGTCTATCGTGTCC[BHQ1]-3’. 5 μl of extracted DNA was used per 20 μl reaction and thermal cycling was performed on an Applied Biosystems 7500 Fast PCR instrument running SDS software v2.3 (ThermoFisher Scientific) under the following conditions: 95 °C for 5 mins, followed by 45 cycles of 95 °C for 10 s and 62 °C for 30 mins.

### Viral entry assays

HFt cells were seeded in 6-well plates at a cell density of 4×10^5^ cells/well 24 h prior to infection. Cells were infected with MPXV or VACV at a MOI 0.1 PFU/cell for 1 h at 37 °C before continued incubation at the desired temperature (as indicated) for a further 1 h. Cell monolayers were washed twice in PBS before being trypsinised. Cells were pelleted at 3000 rpm for 5 mins and resuspended in 200 μl PBS. Total DNA was extracted using QIAamp DNA kit (Qiagen, 51304), according to the manufacturer’s instructions, with a final elution volume of 20 μl. Viral genome levels were quantified by qPCR using virus specific primer probe sets (described above) and normalised to GAPDH using an Applied Biosystems FAM/MGB probe mix (Thermofisher, 4333764F).

### RNA sequencing (RNA-Seq)

HFt cells were seeded onto 12-well plates at a density of 2×10^5^ cells/well. 24 h post-seeding, cells were mock treated or infected with MPXV CVR_S1 (MOI 0.01) for 1h at 37 °C before overlay and continued incubation at 33, 37, 38.5 or 40 °C (as indicated). Cells were washed three times in PBS at 48 h prior to RNA extraction using a RNeasy plus Micro Kit (Qiagen, 74034). Eluted RNA was quantified using Qubit Fluorometer 4 (ThermoFisher, Q33238), Qubit RNA HS Assay (Life Technologies, Q32855) and dsDNA HS Assay Kits (ThermoFisher, Q32854) and quality controlled on a 4200 TapeStation System (Agilent Technologies, G2991A) with a High Sensitivity RNA Screen Tape assay (Agilent Technologies, 5067-5579). All samples had a RIN score of ≥ 8.8. 220 ng of total RNA was used to prepare libraries for sequencing using an Illumina TruSeq Stranded mRNA Library Prep kit (Illumina, 20020594) and SuperScript II Reverse Transcriptase (Invitrogen, 18064014) according to the manufacturer’s instructions. Dual indexed libraries were PCR amplified, purified using AgencourtAMPure XP magnetic beads (Beckman Coulter, 10136224), quantified using Qubit Fluorometer 4 (TheroFisher, Q33238) and Qubit dsDNA HS Assay Kit (ThermoFisher, Q32854), and the size distribution assessed using a 4200 TapeStation System (Agilent Technologies, G2991A) with a High Sensitivity D1000 Screen Tape assay (Agilent Technologies, 5067-5584). Libraries were pooled in equimolar concentrations and sequenced using an Illumina NextSeq 500/550 sequencer (Illumina, FC-404-2005). At least 95% of the reads generated presented a Q score of ≥ 30. RNA-Seq reads were quality assessed using FastQC (http://www.bioinformatics.babraham.ac.uk/projects/fastqc). Sequence adaptors removed using TrimGalore (https://www.bioinformatics.babraham.ac.uk/projects/trim_galore/). RNA-Seq reads were aligned to the *Homo sapiens* genome (GRCh38), downloaded via Ensembl using HISAT2. HISAT2 is a fast and sensitive splice aware mapper, which aligns RNA sequencing reads to mammalian-sized genomes using FM index strategy^67^. FeatureCounts^68^ was used to calculate mapped read counts that were normalized to counts per million (CPM; unless otherwise stated). Generalized linear models (GLMs) with multi-factor designs was used for differential gene expression (DEG) analysis in EdgeR^69^. An FDR (False Discovery Rate) value <0.05 was used as a cut-off of significant differential gene expression. Viral sequences were aligned to the MPXV Clade IIa reference sequence ([NC_063383.1](https://www.ncbi.nlm.nih.gov/nuccore/NC_063383)). High confidence (FDR <0.05, ≥ 1.5 log2 fold change [FC]) DEGs were used for pathway analysis in Reactome (https://reactome.org)^70,71^ or differential pathway analysis in Metascape (https://metascape.org/gp/index.html#/main/step1)^72^. For Reactome analysis, the gene mapping tool was used as a filter to identify pathways enriched (over-represented) for mapped entities. FDR values <0.05 were considered significant for pathway enrichment. In Metascape, all DEGs were used for differential pathway analysis. Pathway *p*-values <0.05 were considered significant. Heat maps were plotted in GraphPad Prism (version 10). Mean counts per million (CPM) values of zero were normalized to 0.01 for log2 presentation. Venn diagrams were plotted using http://bioinformatics.psb.ugent.be/webtools/Venn/. RNA-Seq datasets generated from this study have been deposited in the European Nucleotide Archive (ENA), accession number PRJEB66116.

### Statistical analysis

The number (N) of independent biological experiments is shown throughout. GraphPad Prism (version 10) was used for PCA and statistical analysis. Statistical tests and *p*-values are shown throughout. Statistically significant differences were accepted at *p*<0.05.

## Data availability

All datasets generated and analysed in this study will be made freely available in supplemental files or online (ENA; accession number PRJEB66116) upon manuscript acceptance for publication.

## Acknowledgements

The authors would like to thank the members of the MPOX Research Consortium UK for the constructive comments and feedback during the preparation of this manuscript. This manuscript is dedicated in memory of Kyriaki Nomikou, an exceptional scientist who sadly passed away during the preparation of this manuscript. Kiki was greatly loved and will be sadly missed by everyone who was blessed to meet her.

## Author Contributions

**Data curation:** Chris Davis, Ilaria Epifano, Kieran Dee, Steven McFarlane, Quan Gu, Vattipally B. Sreenu, Lily Tong, Kyriaki Nomikou, Lauren Orr, Chris Boutell.

**Formal analysis:** Chris Davis, Ilaria Epifano, Kieran Dee, Joanna K. Wojtus, Quan Gu, Vattipally B. Sreenu, Chris Boutell.

**Funding acquisition:** Benjamin Brennan, Ana Da Silva Filipe, Emma Thomson, Chris Boutell.

**Investigation:** Chris Davis, Ilaria Epifano, Kieran Dee, Lauren Orr, Antonia Ho, Chris Boutell.

**Methodology:** Chris Davis, Ilaria Epifano, Steven McFarlane, Benjamin Brennan, Lauren Orr, Chris Boutell.

**Project administration:** Ana Da Silva Filipe, Antonia Ho, David. A. Barr, Emma Thomson, Chris Boutell.

**Resources:** Steven McFarlane, Benjamin Brennan, Ana Da Silva Filipe, Antonia Ho, David. A. Barr, Emma Thomson, Chris Boutell.

**Supervision:** Ana Da Silva Filipe, Emma Thomson, Chris Boutell.

**Validation:** Chris Davis, Ilaria Epifano, Kieran Dee, Chris Boutell.

**Visualization:** Chris Davis, Ilaria Epifano, Chris Boutell.

**Writing – original draft:** Chris Davis, Chris Boutell.

**Writing – review & editing:** Chris Davis, Ilaria Epifano, Chris Boutell.

---

**Fig S1.**
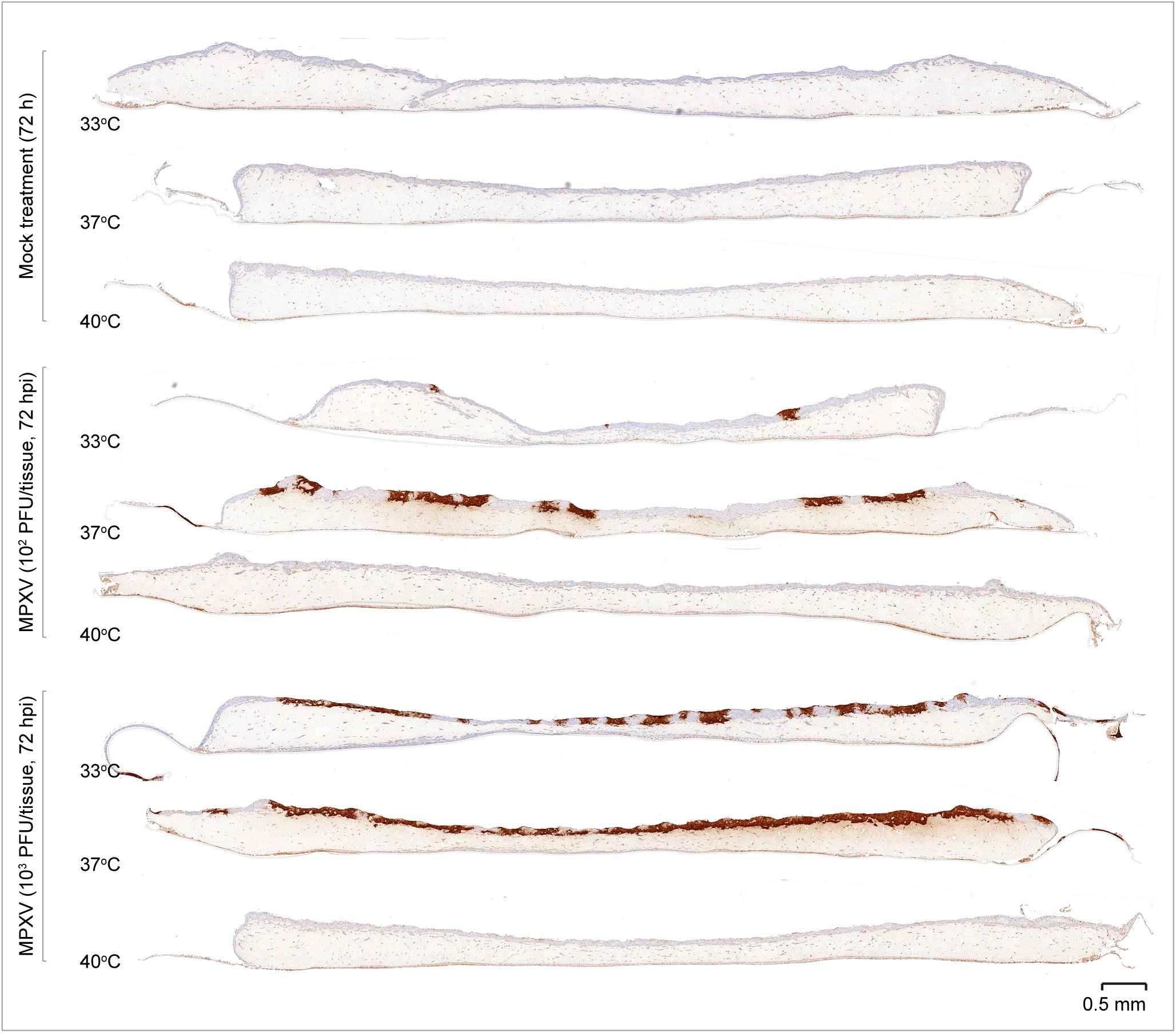
Tissue temperature influences the outcome of MPXV replication in skin epithelium. Human keratinocytes were seed onto fibroblast-containing collagen coated 6.5 mm transwells and differentiated under ALI for 14 days. Pseudostratified skin epithelium cultures were mock treated (media only) or MPXV infected (MOI 10^2^ or 10^3^ PFU/tissue) for 1 h at 37 °C prior to incubation at 33, 37, or 40 °C (as indicated) for 72 h. Whole tissue sections were stained for MPXV virion antigen expression by immunohistochemistry and counter stained with eosin. Scale bar = 0.5 mm.

**Fig S2.**
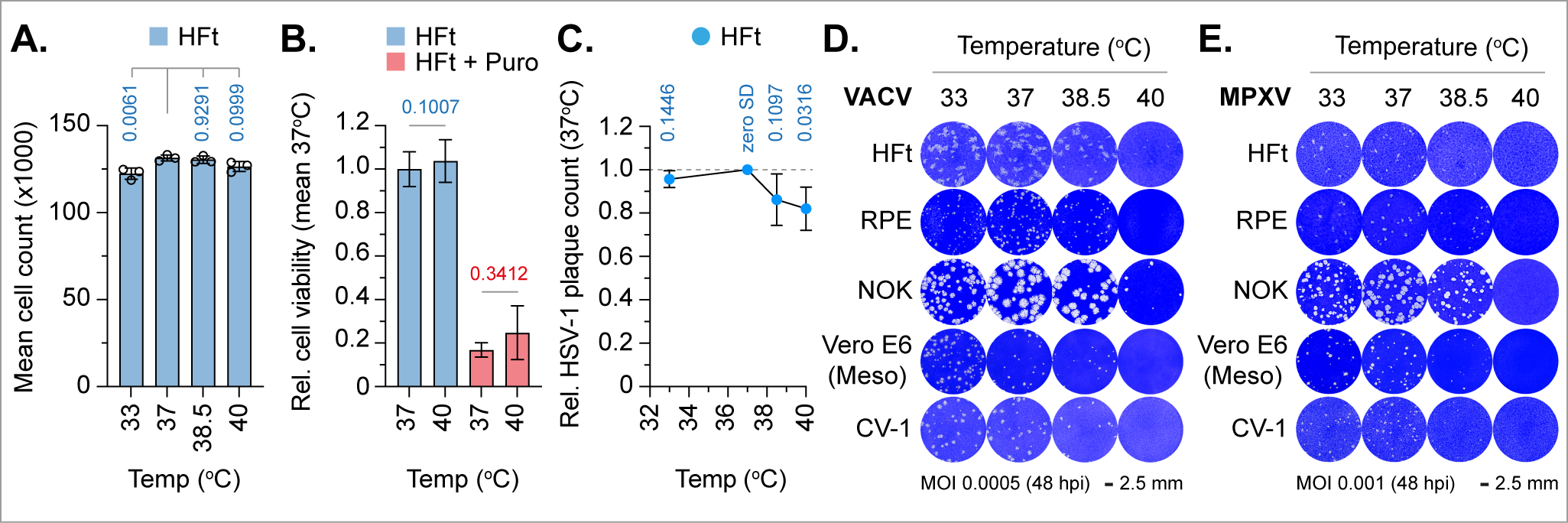
Temperature elevation inhibits OPXV replication. (A) Quantitation of DAPI stained nuclei in mock treated HFt cells incubated at 33, 37, 38.5 or 40 °C for 48 h. (B) MTS cell viability assay of mock treated or puromycin (Puro, 1 μg/ml; positive death control) treated HFt cells incubated at 37 or 40 °C for 24 h (as indicated). (C to E) Confluent HFt (human foreskin fibroblast), NOK (human normal oral keratinocyte), RPE (human retinal pigmented epithelial), Vero E6 (green monkey kidney epithelial), or CV-1 (green monkey kidney fibroblast) cell monolayers were infected with (C) HSV-1 (MOI 0.002 PFU/cell), (D) VACV (MOI 0.0005 PFU/cell), or (E) MPXV (MOI 0.001 PFU/cell) for 1 h at 37 °C prior to overlay and incubation at 33, 37, 38.5 or 40 °C. Infected cell monolayers were fixed at 48 h and stained with Coomassie Brilliant blue. (C) Quantitation of HSV-1 plaque counts in HFt cells over the temperature incubation range of analysis. Values were normalised to plaque counts at 37 °C. Means and SD shown; *p*-values shown, one sample two-tailed t test against a theoretical mean of 1. (D/E) Representative images of VACV or MPXV infected HFT, RPE, NOK, Vero E6 (Meso), and CV-1 cell monolayers stained with Coomassie Brilliant blue at 48 h post-infection (hpi). (A to E) *N*≥3 independent biological experiments. Raw values presented in supplemental S9 data.

**Fig S3.**
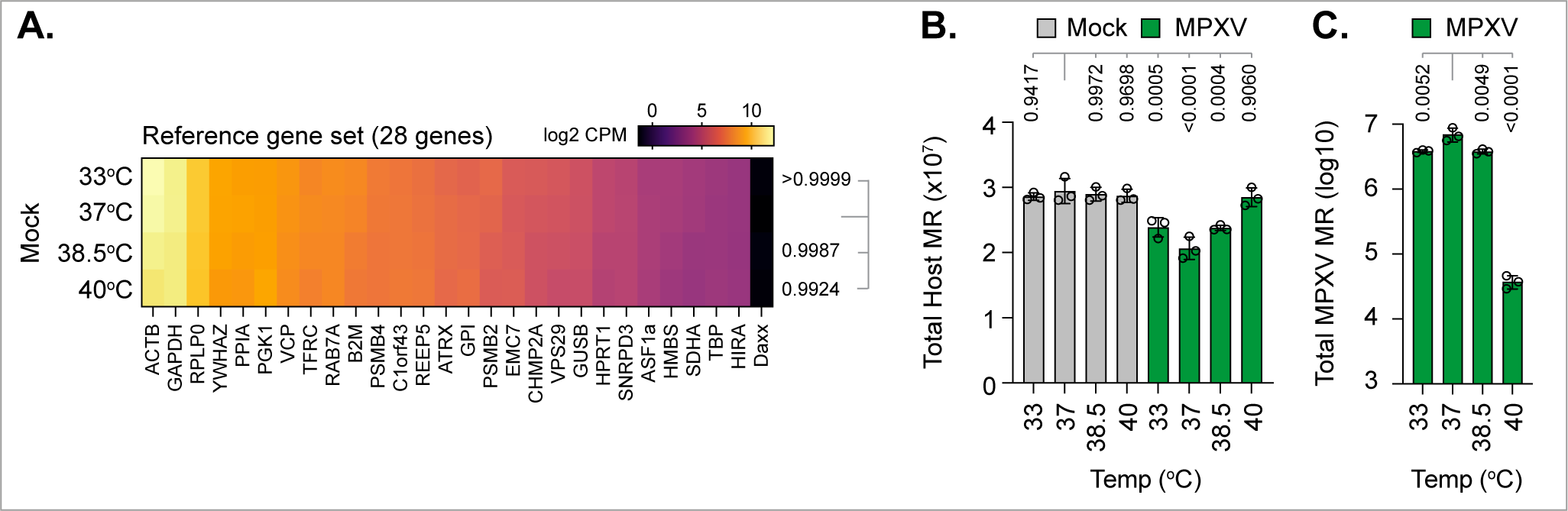
Basal host-cell temperature differentially influences the regulation of MPXV transcription. HFt cells were infected with MPXV (MOI 0.01 PFU/cell) for 1 h at 37 °C prior to incubation at 33, 37, 38.5 or 40 °C. RNA was extracted at 48 h post-infection (hpi) for RNA-Seq. Host sequences were aligned to the human genome and mapped read (MR) counts normalised to counts per million (CPM). Viral sequences were aligned to the MPXV reference sequence ([NC_063383.1](https://www.ncbi.nlm.nih.gov/nuccore/NC_063383)) and viral MR counts used to quantify MPXV transcript abundance. (A) Expression level (log2 CPM) of 28 host reference genes over the temperature range of analysis. (B) Host MR counts in mock treated (grey bars) and MPXV infected (green bars) cells over the temperature range of analysis. (C) Viral MR counts in MPXV infected cells over the temperature range of analysis. (A to C) *N*=3 independent biological experiments; *p*-values shown, unpaired Dunnett’s one-way ANOVA. Raw values presented in supplemental S10 data.

**Fig S4.**
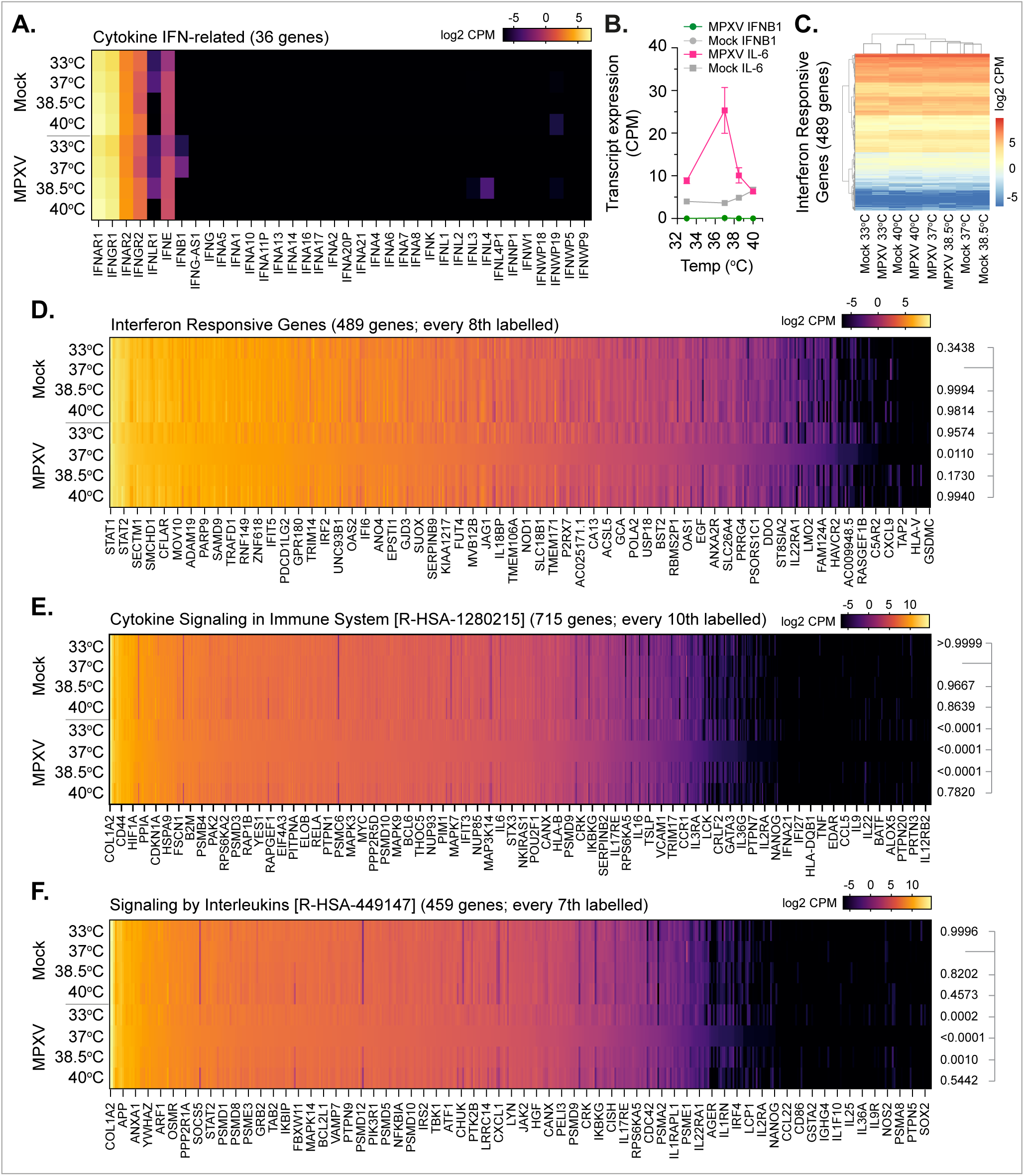
MPXV infection does not stimulate the induction of the type-I IFN response. HFt cells were mock treated or infected with MPXV (MOI 0.01 PFU/cell) for 1 h at 37 °C prior to incubation at 33, 37, 38.5 or 40 °C. RNA was extracted at 48 h post-infection (hpi) for RNA-Seq. Host sequences were aligned to the human genome and mapped read (MR) counts normalised to counts per million (CPM). (A) Expression level (log2 CPM) of host interferon (IFN)-related receptors and cytokines in mock treated or MPXV infected cells over the temperature range of analysis. (B) Expression profile (CPM) of IFNB1 (circles) and IL-6 (squares) transcript levels in mock treated (grey lines) or MPXV infected (pink lines) cells over the temperature range of analysis. Means and SD shown. (C/D) Expression profile (log2 CPM) of 489 interferon responsive genes previously identified to be upregulated in response to universal interferon^73^; every 8^th^ gene labelled. Grey lines in C highlight profile similarity between experimental conditions. (E) Expression profile (log2 CPM) of host genes associated with Cytokine Signalling in Immune System (R-HAS-1280215; 715 genes in total) in mock treated or MPXV infected cells over the temperature range of analysis. (F) Expression profile (log2 CPM) of host genes associated with Signalling by Interleukins (R-HAS-449147; 459 genes in total) in mock treated or MPXV infected cells over the temperature range of analysis. (A to G) *N*=3 independent biological experiments. Raw values presented in supplemental S11 data. (D to F) *P*-values shown, Dunnett’s paired one-way ANOVA with Giesser-Greenhouse correction.

**Fig S5.**
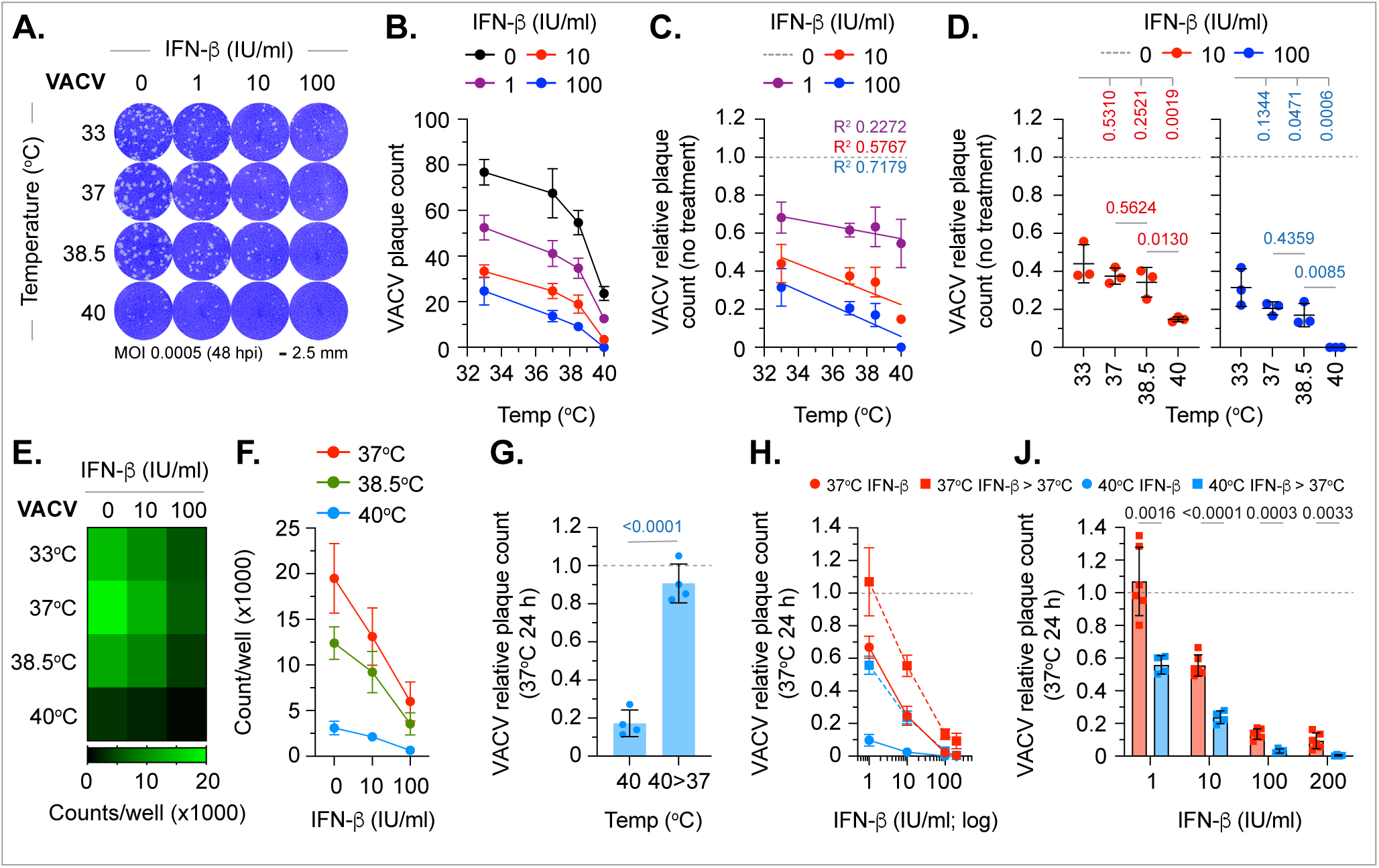
Temperature elevation enhances the type-I IFN-mediated host-cell restriction of VACV. (A to D) HFt cells were pre-treated with IFN-ϕ3 (0 to 100 IU/ml, as indicated) for 16 h at 33, 37, 38.5 or 40 °C prior to VACV (MOI 0.0005 PFU/cell) infection (1 h at 37 °C) and continued incubation at their respective temperatures in the presence of IFN. (A) Representative images of VACV infected cell monolayers pre-treated with IFN and stained with Coomassie Brilliant blue at 48 h post-infection (hpi). (B) Quantitation of VACV plaque counts at 48 hpi (as shown in A). Means and SD shown. (C/D) Relative VACV plaque counts). Values were normalised to no IFN treatment (dotted grey line) per incubation temperature. (C) Means and SD shown; coloured lines and text, linear regression and corresponding R^2^ values. (D) As in C, all data points shown; black line, mean; whisker, SD; *p*-values shown, Dunnett’s unpaired one-way ANOVA. (E to J) Naïve HFt cells were infected with VACV (MOI 0.01 PFU/cell, E and F; MOI 0.0005 PFU/cell, G to J) for 1 h at 37 °C prior to overlay with media containing IFN-β (0 to 200 IU/ml, as indicated) and incubation at 33, 37, 38.5 or 40 °C. (E/F) Cells were fixed at 24 h and stained for VACV virion protein expression and the number of antigen positive cells quantified by indirect immunofluorescence. (E) Mean VACV positive cell counts (x1000) per infected cell monolayer. (F) As in E, mean and SD shown. (G to J) Infected or infected and IFN treated monolayers incubated at 37 or 40 °C were either fixed at 24 h or washed and overlayed with fresh media (IFN washout) and incubation at 37 °C for an additional 24 h prior to fixation and staining. (G) Quantitation of VACV plaque counts in cell monolayers incubated at 40 °C or temperature downshifted from 40 to 37 °C (40>37) with continued incubation for 24 h. (H) Quantitation of VACV plaque counts in cell monolayers incubated at 37 or 40 °C in the presence of IFN (red and blues circles plus solid lines, respectively) or following IFN washout and continued incubation at 37 °C for 24 h (red and blue squares plus dotted lines, respectively). (J) Values presented for IFN washout and continued incubation at 37 °C for 24 (red and blue squares in H), all data points shown. (G to J) Values were normalised to plaque counts determined at 37 °C (no IFN treatment at 24 h). Mean and SD shown; *p*-values shown, unpaired two-tailed *t* test. Raw values presented in supplemental S12 Data.

